# NHR-49/PPAR-α and HLH-30/TFEB promote *C. elegans* host defense via a flavin-containing monooxygenase

**DOI:** 10.1101/2020.09.03.282087

**Authors:** Khursheed A. Wani, Debanjan Goswamy, Stefan Taubert, Ramesh Ratnappan, Arjumand Ghazi, Javier E. Irazoqui

## Abstract

During bacterial infection, the host is confronted with multiple overlapping signals that are integrated at the organismal level to produce defensive host responses. How multiple infection signals are sensed by the host and how they elicit the transcription of host defense genes is much less understood at the whole-animal level than at the cellular level. The model organism *Caenorhabditis elegans* is known to mount transcriptional defense responses against intestinal bacterial infections that elicit overlapping starvation and infection responses, but the regulation of such responses is not well understood. Direct comparison of *C. elegans* that were starved or infected with *Staphylococcus aureus* revealed a large infection-specific transcriptional signature. This signature was almost completely abrogated by deletion of transcription factor *hlh-30/TFEB*, except for six genes including a flavin-containing monooxygenase (FMO) gene, *fmo-2/FMO5*. Deletion of *fmo-2/FMO5* severely compromised infection survival, thus identifying the first FMO with innate immunity functions in animals. Moreover, the mechanism of *fmo-2/FMO5* induction required the nuclear hormone receptor, NHR-49/PPAR-α, which induced *fmo-2/FMO5* and host defense cell non-autonomously. These findings for the first time reveal an infection-specific host response to *S. aureus*, identify HLH-30/TFEB as its main regulator, reveal that FMOs are important innate immunity effectors in animals, and identify the mechanism of FMO regulation through NHR-49/PPAR-α in *C. elegans*, with important implications for innate host defense in higher organisms.

## INTRODUCTION

In their natural habitat, *C. elegans* feed on microbes that grow on rotting vegetable matter, and thus face a high likelihood of ingesting pathogens (Schulenburg and Felix, 2017). To defend against infection, *C. elegans* possess innate host defense mechanisms that promote their survival (Ermolaeva and Schumacher, 2014; Kim and Ewbank, 2018). In the laboratory, model human pathogenic bacteria cause intestinal pathology and death through poorly understood mechanisms (Irazoqui et al., 2010a). Infected animals experience both chemical signals that reveal the pathogen’s presence and organismal stress caused by the infection. Over the last 15 years, several studies have identified and characterized *C. elegans* gene expression changes in response to pathogenic bacteria, fungi, and viruses, mounted through evolutionarily conserved mechanisms (Irazoqui et al., 2010b; Kim and Ewbank, 2018). However, the relative contributions of pathogen sensing and organismal stress mechanisms to the total pathogen-induced response remain unclear.

We previously showed that ingested Gram-positive bacterium *Staphylococcus aureus* causes drastic cytopathology in *C. elegans* (Irazoqui et al., 2010a). Infection with *S. aureus* results in progressive effacement and lysis of intestinal epithelial cells, whole-body cellular breakdown, and death (Irazoqui et al., 2010a). Therefore, *S. aureus*-infected *C. elegans* experience dietary changes from its laboratory food of nonpathogenic *E. coli*, as well as intestinal destruction, cellular stress, and putative molecular signals produced by the pathogen.

In previous work, we showed that *C. elegans* mount a pathogen-specific transcriptional host response against *S. aureus*, which includes genes that encode antimicrobial proteins (*e.g*. lysozymes, antimicrobial peptides, and secreted C-type lectins) and cytoprotective factors (*e.g*. autophagy genes, lysosomal factors, and chaperones) that are necessary and sufficient for survival (Irazoqui et al., 2010a). However, the relative contributions of organismal stress and pathogen detection to the induction of the overall host defense response are unknown.

We recently discovered that the induction of a large majority of the transcriptional host response to *S. aureus* requires HLH-30, the *C. elegans* homolog of mammalian transcription factor EB (TFEB) (Visvikis et al., 2014). TFEB belongs to the MiT family of transcription factors, which in mammals and *C. elegans* controls the transcription of autophagy and lysosomal genes in response to nutritional stress in addition to infection (Lapierre et al., 2013; Raben and Puertollano, 2016). HLH-30 and TFEB also regulate lipid store mobilization during nutritional deprivation (O’Rourke and Ruvkun, 2013; Settembre et al., 2013). Thus, HLH-30/TFEB could potentially integrate organismal stress, metabolism, and pathogen recognition to elicit coordinated host responses to infection. How HLH-30/TFEB integrates this information to produce stress-specific responses and what other factors are involved in such specificity are poorly understood. Specifically, the genes that are induced during infection independently of nutritional stress are not known.

Here we report that *S. aureus* infection in *C. elegans* elicits a transcriptional response that is distinct from that induced by nutritional deprivation, thus defining an infection-specific transcriptional signature. Both the starvation response and the infection-specific signature were largely dependent on HLH-30/TFEB, highlighting its key role as a transcriptional integrator of organismal stress during infection. Moreover, we identified six genes that were specifically induced during infection even in the absence of HLH-30/TFEB, potentially revealing an alternative transcriptional host response signaling pathway. The induction of one of the six genes, *fmo-2/FMO5*, was entirely and non cell-autonomously dependent on transcription factor NHR-49/PPAR-α (Van Gilst et al., 2005), suggesting that NHR-49/PPAR-α defines a novel host infection defense pathway. Moreover, functional characterization of *fmo-2/FMO5* suggested that its enzymatic activity is specifically required for host defense against *S. aureus*, revealing that FMO-2/FMO5 is a key host defense effector. In addition to identifying a new transcriptional regulator of the host defense response, this is the first report that shows that flavin-containing monooxygenases such as FMO-2/FMO5 are important for host defense in animals.

## RESULTS

### Starvation and infection trigger distinct transcriptional responses

Our prior studies showed that *S. aureus* infection of *C. elegans* causes a robust host transcriptional response that results in the upregulation of 825 genes (Irazoqui et al., 2010a). It is likely that this transcriptional response to infection is compounded with nutritional stress, due to nutritional differences between laboratory food nonpathogenic *E. coli* and *S. aureus*, and due to intestinal destruction caused by the pathogen (Irazoqui et al., 2010a). To identify genes that are induced during infection independently of nutritional stress, we used whole-animal RNA-seq to directly compare infected and starved animals (**Fig. 1A**). We identified 388 genes that were differentially expressed between these two conditions (**Fig. 1B, C, Table S1**). About 70% (283 genes) of differentially expressed genes were upregulated by starvation, while about 30% (105 genes) were upregulated by infection (**Table S1)**. Gene ontology analysis showed the starvation-induced genes to belong mostly to metabolic processes, whereas the infection-specific signature was highly enriched for innate immune response genes (**Table S2**). RT-qPCR of the 13 most highly infection-induced genes relative to animals that were starved or fed nonpathogenic *E. coli* laboratory food confirmed their *S. aureus-*specific induction (**Fig. 1D, Fig. S1A**). Thus, we identified an infection-specific signature of genes that excludes expression changes that are caused by starvation, indicating that the host responses to nutritional deprivation and *S. aureus* infection have distinct and specific features.

**Figure 1.**
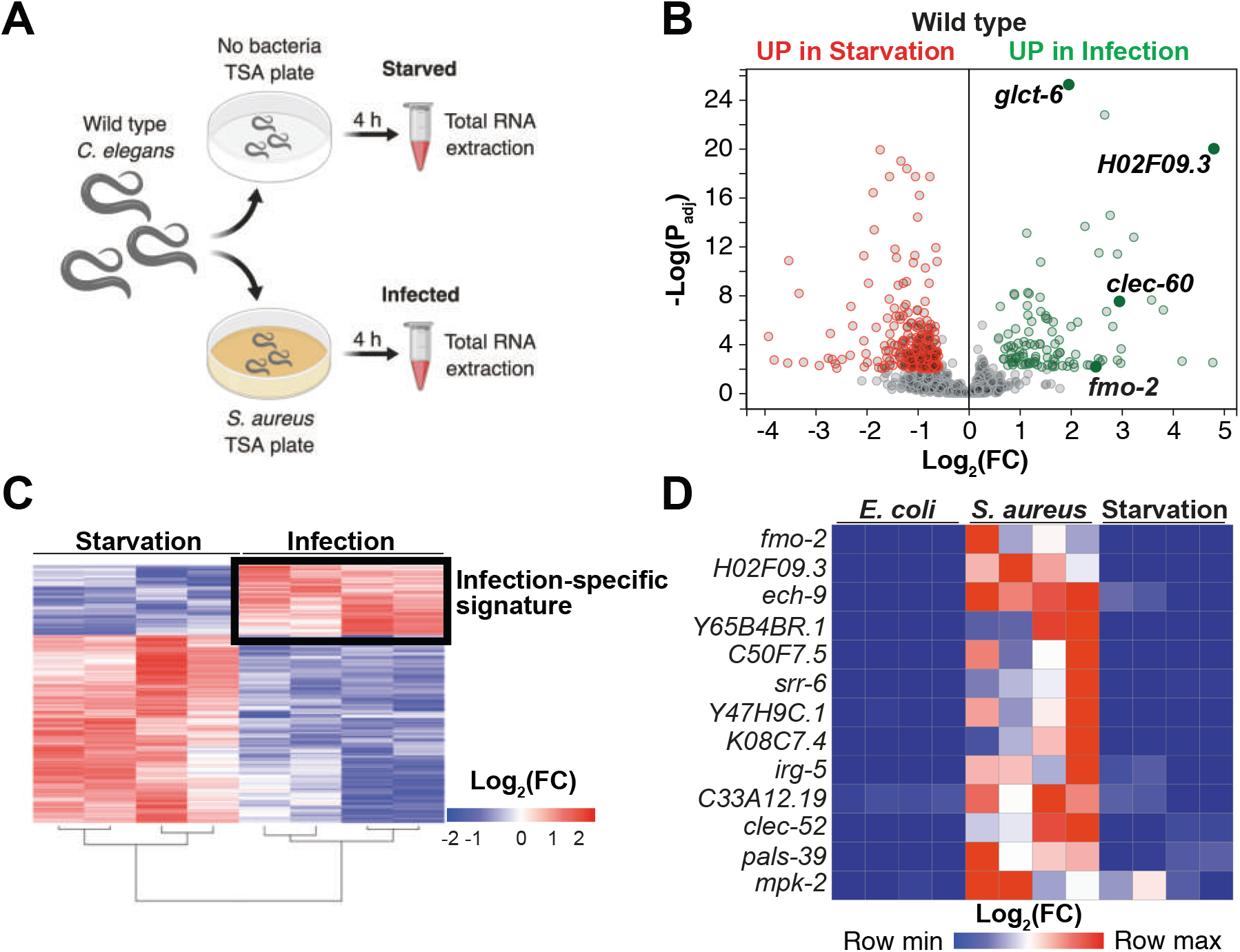
Starvation and *S. aureus* infection trigger distinct transcriptional responses. **(A)** Schematic overview of experimental approach for RNA-seq conditions. Synchronized young adults were subjected to either starvation or infection for 4 h before RNA extraction. **(B)** Volcano plot of differentially expressed genes (P_adj_ ≤ 0.01). Genes that were induced in each condition relative to the other are indicated in red (for starvation) and green (for infection). FC, fold change. P_adj_, adjusted P value. **(C)** Heat map of differentially expressed genes [Log_2_(FC)] comparing infection with *S. aureus* SH1000 to starvation by RNA-seq. The boxed area represents the designated infection-specific expression signature. **(D)** Heat map of a set of 13 genes most highly induced by *S. aureus* SH1000 compared to starvation, whose relative transcript levels were measured by RT-qPCR and plotted as row-normalized log2(relative expression), or -ΔCt. Conditions include nonpathogenic *E. coli, S. aureus* (4 h), and starvation (4 h). Columns represent independent biological replicates.

### HLH-30/TFEB is critical for host responses to starvation and infection

HLH-30/TFEB was shown to be important for gene induction during dietary challenge and during infection (O’Rourke and Ruvkun, 2013; Settembre et al., 2013; Visvikis et al., 2014). However, whether HLH-30/TFEB regulates the infection-specific response was not known. To assess the relevance of HLH-30/TFEB to the infection-specific signature, we compared starved and infected *hlh-30/TFEB* loss of function mutants by RNA-seq. To our surprise, in *hlh-30/TFEB* mutants differential gene expression between starvation and infection was almost completely abrogated (**Fig. 2A, Table S3**). Of the 105 genes in the infection-specific signature, only 6 were induced in *hlh-30/TFEB* mutants (**Fig. 2B**), including *clec-52, fmo-2/FMO5*, and the uncharacterized genes *C33A12.19, C54F6.12, K08C7.4*, and *Y47H9C.1* (**Table S3**). RT-qPCR confirmed the predicted results for the selected 13 top induced genes (**Fig. 2C, Figure S1B**). Particularly, we verified that *fmo-2/FMO5* was partially induced in *hlh-30/TFEB* mutants compared to wild type. Partial induction of *fmo-2/FMO5* in *hlh-30/TFEB* mutants was rescued by transgenic re-expression of *hlh-30/TFEB* driven by its endogenous promoter (**Fig. 2D**). Altogether, these results showed that HLH-30/TFEB is crucial for both the starvation and the infection-specific responses, and hinted at an HLH-30/TFEB-independent pathway for the induction of 6 infection-specific genes.

**Figure 2.**
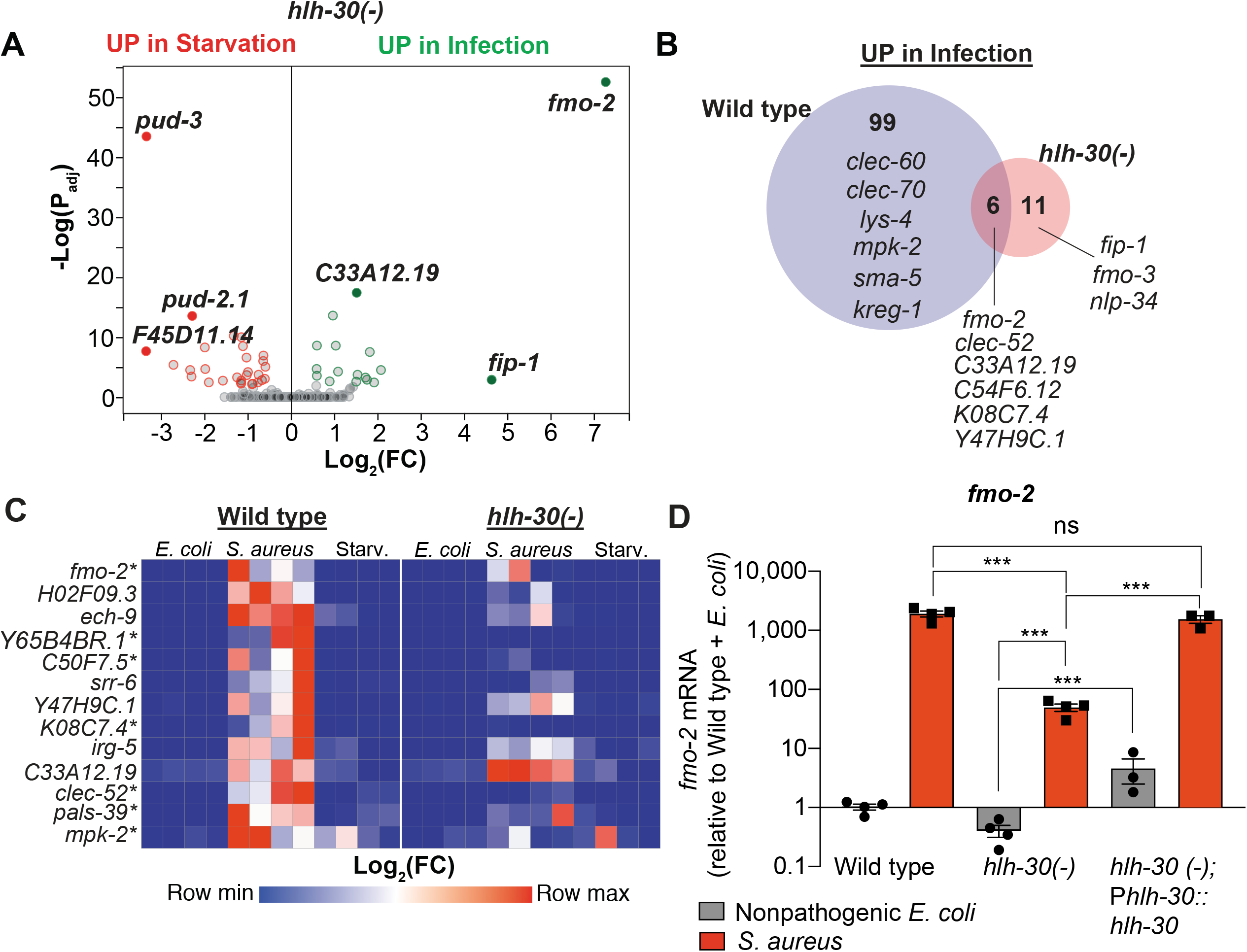
HLH-30/TFEB is critical for host responses to starvation and infection. **(A)** Volcano plot of differentially expressed genes in *hlh-30/TFEB* loss of function mutants (P ≤ 0.01). Genes that were induced in each condition relative to the other are indicated in red (for starvation) and green (for infection). **(B)** Venn diagram representing genes that were upregulated during infection compared to starvation in wild type and *hlh-30/TFEB* mutants. A few selected genes are indicated for reference. **(C)** Heat map of RT-qPCR (-ΔCt) relative expression values of a set of 13 genes most highly induced by *S. aureus* v starvation in wild type, measured in wild type and *hlh-30/TFEB* mutants. Conditions include nonpathogenic *E. coli, S. aureus*, and starvation. Columns represent independent biological replicates. * indicates genes that were highly induced in wild type compared to *hlh-30/TFEB* mutants during infection, and thus were partially or completely HLH-30/TFEB-dependent. “Starv.”, starvation. **(D)** RT-qPCR of *fmo-2/FMO5* transcript in wild type, *hlh-30/TFEB* loss of function mutants, and *hlh-30(-); Phlh-30::hlh-30::gfp* (complemented) animals fed nonpathogenic *E. coli* or infected with *S. aureus* (4 h). Data are normalized to wild type fed nonpathogenic *E. coli*, means ± SEM (3 - 4 independent biological replicates). *** P < 0.001, ns = not significant, one-way ANOVA followed by Sidak’s test for multiple comparisons.

### Infection induces *fmo-2/FMO5* via NHR-49/PPAR-α

As the most highly induced infection-specific gene in *hlh-30/TFEB* mutants (**Fig. 2A, Table S3**), *fmo-2/FMO5* attracted our attention. Our prior studies showed that infection causes *fmo-2/FMO5* induction independently of previously identified host defense pathways, including p38 MAPK, TGF-β, ERK, insulin, Wnt, and HIF-1 pathways (Irazoqui et al., 2008, 2010a; Luhachack et al., 2012; Visvikis et al., 2014). Additionally, we found that *fmo-2/FMO5* can be partially induced independently of HLH-30/TFEB (**Fig. 2** and (Visvikis et al., 2014)). Therefore, additional transcriptional regulators must be involved in *fmo-2/FMO5* induction.

Previous studies identified NHR-49, a nuclear receptor homologous to human PPAR-α and HNF4-α, as essential for *fmo-2/FMO5* induction during exogenous oxidative stress (Goh et al., 2018; Hu et al., 2018). To examine the role of NHR-49/PPAR-α during infection, we checked *fmo-2/FMO5* expression in *nhr-49/PPARA* null mutants (Liu et al., 1999; Van Gilst et al., 2005). We found that in *nhr-49/PPARA* null mutants, expression of the *fmo-2/FMO5* fluorescent transcriptional reporter was decreased both under normal conditions (**Fig. 3E**) and, importantly, was barely induced by *S. aureus* in the intestinal epithelium (**Fig. 3F**). In contrast, *fmo-2/FMO5* induction by *S. aureus* was partially dependent on *hlh-30/TFEB*, as predicted by RNA-seq (**Fig. 3A-D, Fig. 2**); in *hlh-30/TFEB* mutants expression was preserved in the pharyngeal isthmus, pharyngeal-intestinal valve, nervous system, coelomocytes, and posterior intestinal epithelium (**Fig. 3D**). Thus, we found that HLH-30/TFEB appears to contribute to *fmo-2/FMO5* transcription in the intestinal epithelium, while *nhr-49/PPARA* appears to be more important in other tissues. By RT-qPCR, noninfected *nhr-49/PPARA* mutants exhibited about 10-fold lower *fmo-2/FMO5* expression than wild type (**Fig. 3G**). After infection, *nhr-49/PPARA* mutants completely failed to induce *fmo-2/FMO5* (**Fig. 3G**). Moreover, transgenic rescue of *nhr-49/PPARA* driven by its endogenous promoter partially restored *fmo-2/FMO5* induction (**Fig. 3G**). RT-qPCR of the other five HLH-30-independent genes showed that *K08C7.4* induction was also dependent on NHR-49/PPAR-α (**Fig. S2C**). These data suggested that NHR-49/PPAR-α contributes to the induction of some of the HLH-30-independent host defense genes, but the biological significance of NHR-49/PPAR-α to host defense was not clear.

**Figure 3.**
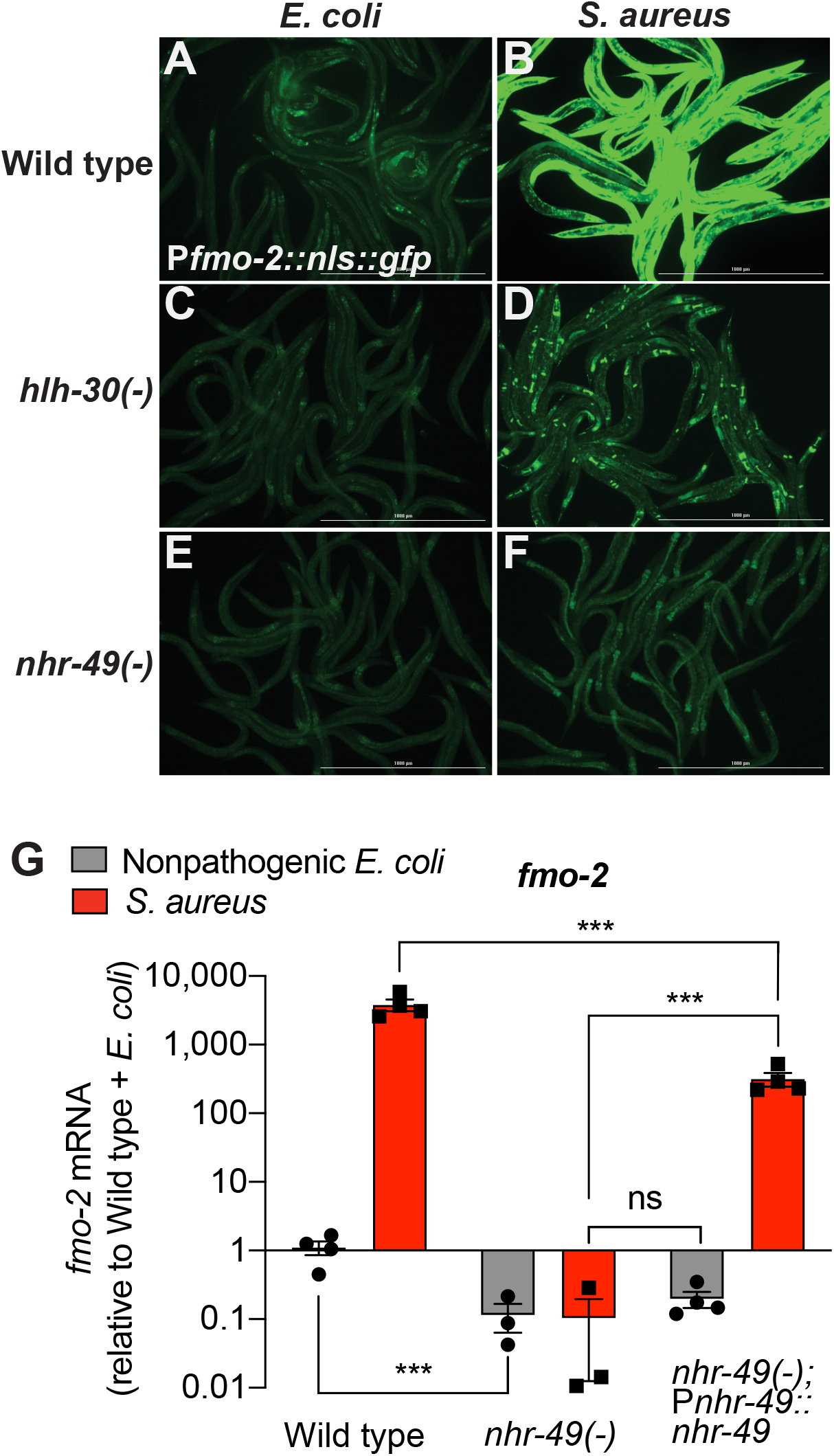
Infection induces *fmo-2/FMO5* via NHR-49/PPAR-α. **(A-F)** Epifluorescence micrographs of animals carrying *Pfmo-2::nls::gfp* in wild type (**A, B**), *hlh-30/TFEB* **(C, D)**, and *nhr-49/PPARA* mutant backgrounds **(E, F)** after feeding on *E. coli* OP50 or infection with *S. aureus* SH1000 (4 h). Scale bar = 1,000 μm. **(C)** Relative expression of *fmo-2/FMO5* transcript (RT-qPCR -ΔCt) in wild type, *nhr-49/PPARA* loss of function mutants, and *nhr-49(-); Pnhr-49::nhr-49* (complemented) animals fed nonpathogenic *E. coli* OP50 or infected with *S. aureus* SH1000 (4 h). Data are normalized to wild type on *E. coli*, means ± SEM (3 - 4 independent biological replicates). *** P < 0.001, ns = not significant, one-way ANOVA followed by Sidak’s test for multiple comparisons.

### NHR-49/PPAR-α is required for host defense

Compared to wild type, null *nhr-49/PPARA* mutants showed defective survival of *S. aureus* infection (**Fig. 4A**) and shorter lifespan when fed nonpathogenic *E. coli* (**Fig. 4B**), as previously reported (Van Gilst et al., 2005). These results suggested that NHR-49/PPAR-α may have important roles in both host defense and aging. Transgenic rescue of *nhr-49/PPARA* driven by its endogenous promoter completely rescued the infection survival defect (**Fig. 4A**) but only partially restored the total lifespan on *E. coli* (**Fig. 4B**), suggesting that distinct thresholds of NHR-49/PPAR-α function exist in infection and aging.

**Figure 4.**
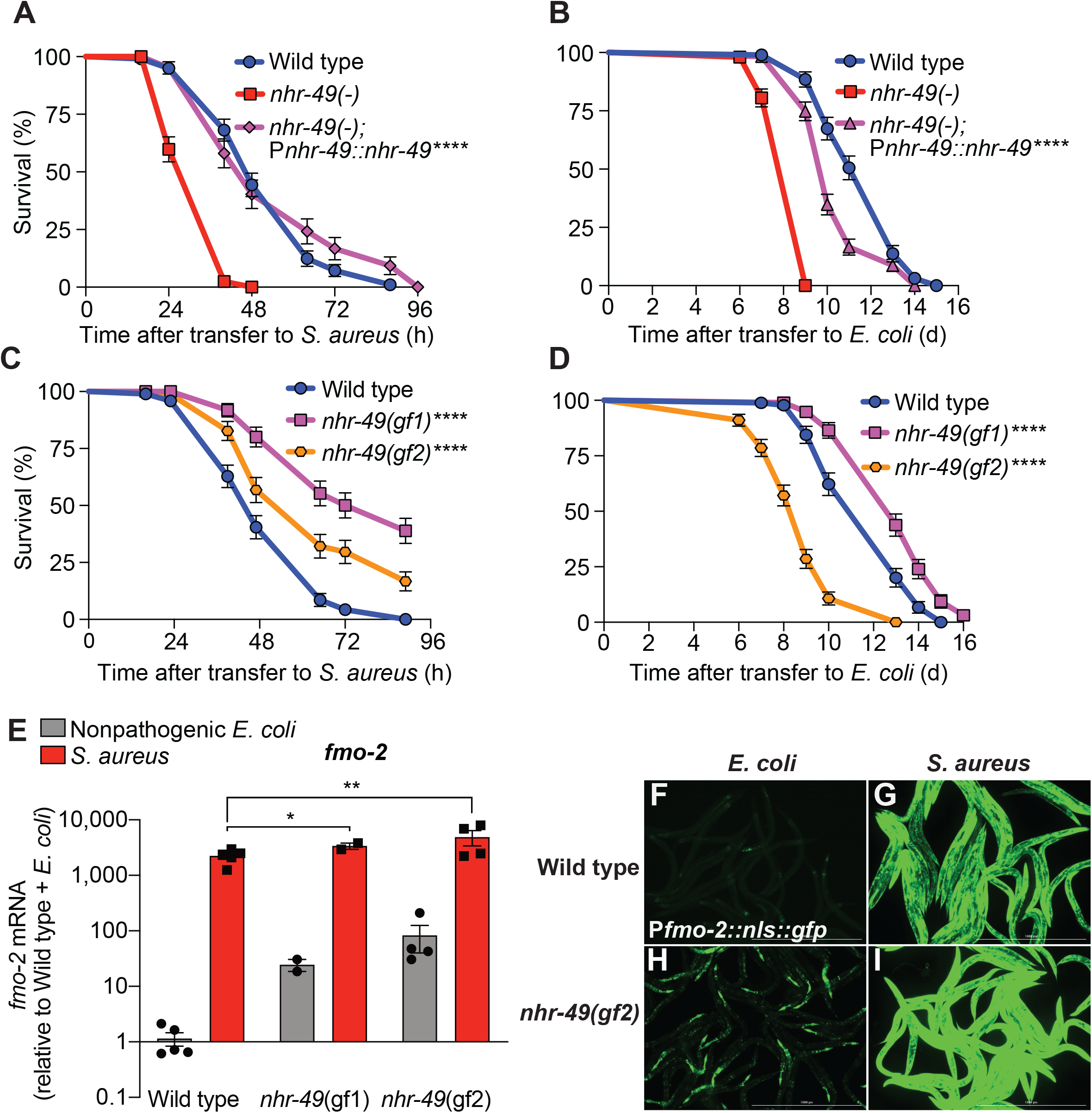
NHR-49/PPAR-α is required for host defense against infection. **(A)** Survival of wild type, *nhr-49/PPARA* loss of function, and *nhr-49(-); Pnhr-49::nhr-49* (complemented) animals infected with *S. aureus* SH1000. Data are representative of 2 independent replicates. **** P < 0.0001 (Log-Rank test). **(B)** Lifespan on nonpathogenic *E. coli* OP50 of wild type, *nhr-49/PPARA* loss of function, and *nhr-49(-); Pnhr-49::nhr-49* animals. Data are representative of 2 independent replicates. **** P < 0.0001 (Log-Rank test). **(C)** Survival of wild type and two *nhr-49/PPARA* gain of function mutants (gf1 = et7 and gf2 = et8) infected with *S. aureus* SH1000. Data are representative of 2 independent replicates. **** P < 0.0001 (Log-Rank test). **(D)** Lifespan of wild type and *nhr-49/PPARA* gain of function mutants on *E. coli* OP50. Data are representative of 3 independent replicates. **** P < 0.0001 (Log-Rank test). **(E)** Relative expression of *fmo-2/FMO5* transcript (RT-qPCR -ΔCt) in wild type and *nhr-49/PPARA* gain of function mutants fed nonpathogenic *E. coli* OP50 or infected with *S. aureus* SH1000 (4 h). Data are normalized to wild type on *E. coli*, means ± SEM (2 - 5 independent biological replicates). * P < 0.05, ** P < 0.01, unpaired two-sample two-tailed *t*-test. **(F-I)** Epifluorescence micrographs of *Pfmo-2::nls::gfp* in wild type **(F-G)** and *nhr-49*(*gf2*) mutants **(H-I)** fed nonpathogenic *E. coli* OP50 or infected with *S. aureus* SH1000 (4 h). Scale bar = 1,000 μm.

Moreover, relative to wild type, two distinct *nhr-49/PPARA* gain-of-function mutants (Lee et al., 2016; Svensk et al., 2013) showed enhanced infection survival (**Fig. 4C**). In contrast, compared to wild type, gain-of-function mutant *nhr-49*(*et7*) (gf1) exhibited prolonged lifespan on *E. coli*, while *nhr-49*(*et8*) (gf2) exhibited shortened lifespan (**Fig. 4D**), consistent with previous results (Lee et al., 2016). These results show that NHR-49/PPAR-α promotes host infection survival, while its function in aging may be more complex.

In line with previous results (Lee et al., 2016), RT-qPCR of noninfected animals showed constitutively elevated *fmo-2/FMO5* expression in gain-of-function *nhr-49/PPARA* mutants relative to wild type, consistent with the observed pro-survival function of NHR-49/PPAR-α (**Fig. 4E**). Upon infection, both gain-of-function mutants exhibited further *fmo-2/FMO5* induction, reaching higher *fmo-2/FMO5* expression than wild type controls (**Fig. 4E**). Consistently, on nonpathogenic *E. coli nhr-49/PPARA* gain-of-function mutants exhibited constitutively high *fmo-2/FMO5* reporter GFP expression in the anterior pharynx, pharyngeal isthmus, nervous system, and the anterior intestinal epithelium (**Fig. 4F, H**). Infection further increased reporter expression throughout the body in *nhr-49/PPARA* gain-of-function mutants, becoming much stronger than wild type animals (**Fig. 4G, I**). These results suggested that *nhr-49/PPARA* activation is sufficient for *fmo-2/FMO5* expression in a spatially restricted pattern, and confirmed that infection synergistically upregulates *fmo-2/FMO5* expression throughout the entire body. Interestingly, the pattern of *fmo-2/FMO5* expression in noninfected *nhr-49/PPARA* gain-of-function mutants (**Fig. 4H**) resembled that observed in infected *hlh-30/TFEB* null mutants (**Fig. 3D**), suggesting that *nhr-49/PPARA* may drive *fmo-2/FMO5* expression in the pharynx, nervous system, and anterior intestine, while *hlh-30/TFEB* may do so in the intestine, muscle, and epidermis. Therefore, HLH-30/TFEB and NHR-49/PPAR-α may have complementary roles for the spatial pattern of *fmo-2/FMO5* expression.

### NHR-49/PPAR-α functions in multiple tissues for host defense

*nhr-49/PPARA* is expressed in multiple tissues (Ratnappan et al., 2014). To identify specific tissues where *nhr-49/PPARA* is sufficient for host defense, we reintroduced wild type *nhr-49/PPARA* into *nhr-49/PPARA* mutants driven by tissue-specific promoters, including intestine, neurons, muscle, and epidermis (*i.e*. hypodermis). We examined these rescue lines for *fmo-2/FMO5* induction and survival of infection. Intestinal rescue of *nhr-49/PPARA* fully restored both basal and induced *fmo-2/FMO5* expression (**Fig. 5A**). Re-expression of *nhr-49/PPARA* partially restored *fmo-2/FMO5* induction in other lines (**Fig. 5D, G, J**). Consistently, expression in each of these tissues also rescued the infection survival defect of *nhr-49/PPARA* mutants. In fact, intestinal, neuronal, and muscular expression produced enhanced infection survival compared to wild type (**Fig. 5B, E, H**), while epidermal expression rescued infection survival to a level similar to wild type (**Fig. 5K**). These results suggested that *nhr-49/PPARA* can function from any one of these tissues to promote host defense against infection.

**Figure 5.**
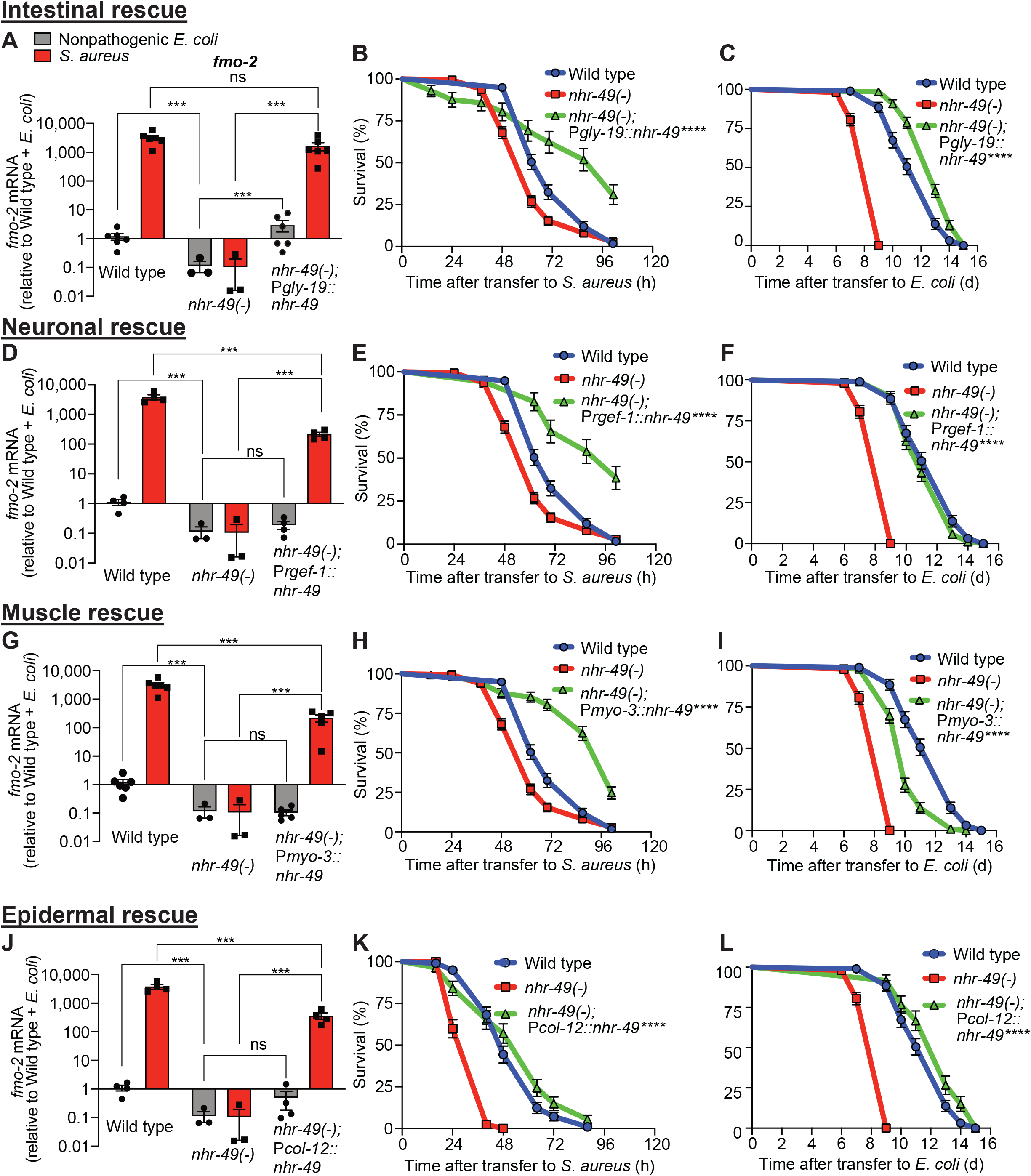
NHR-49/PPAR-α functions in multiple tissues for host defense. **(A, D, G, J)** Relative expression of *fmo-2/FMO5* transcript (RT-qPCR -ΔCt) in wild type, *nhr-49/PPARA* loss of function mutants, and tissue-specific *nhr-49/PPARA* rescue lines: *Pglp-19* for intestine, (*Pglp-19::nhr-49::gfp*), *Prgef-1* for nervous system (*Prgef-1::nhr-49::gfp*), *Pmyo-3* for body wall muscle (*Pmyo-3::nhr-49::gfp*), and *Pcol-12* for epidermis (*Pcol-12::nhr-49::gfp*); fed nonpathogenic *E. coli* OP50 or infected with *S. aureus* SH1000 (4 h). Data are normalized to wild type on *E. coli*, means ± SEM (3 - 6 independent biological replicates,). *** P < 0.001, ns = not significant, one-way ANOVA followed by Sidak’s test for multiple comparisons. **(B, E, H, K)** Survival of wild type, *nhr-49/PPARA* loss of function, and tissue-specific *nhr-49/PPARA* rescue lines infected with *S. aureus*. Data are representative of 2 independent replicates. **** P < 0.0001 (Log-Rank test). Comparisons are made between *nhr-49*(-) and the rescue lines. **(C, F, I, L)** Lifespan of wild type, *nhr-49/PPARA* loss of function, and tissue-specific *nhr-49* rescue lines on nonpathogenic *E. coli*. Data are representative of 3 independent replicates. **** P < 0.0001 (Log-Rank test). Comparisons are made between *nhr-49*(-) and the rescue lines.

In contrast, tissue-specific complementation of *nhr-49/PPARA* had more complex effects on normal lifespan on nonpathogenic *E. coli*. Intestinal and epidermal expression not only rescued *nhr-49/PPARA* mutant lifespan but also prolonged it compared to wild type (**Fig. 5C, L**). Neuronal expression rescued lifespan to wild type level (**Fig. 5F**), and muscle expression caused partial rescue (**Fig. 5I**). Together, these data suggest that *nhr-49/PPARA* may play distinct and tissue-specific roles for infection survival and lifespan.

### NHR-49/PPAR-α controls a fraction of the infection-specific transcriptional signature

To better understand the biological relevance of *nhr-49/PPARA* to the infection-specific host response, we compared the transcriptomes of starved and infected *nhr-49/PPARA* mutants. In stark contrast to *hlh-30/TFEB* mutants, which showed a much-reduced differential response compared to wild type (**Fig. 2**), *nhr-49/PPARA* mutants exhibited many more differentially expressed genes than wild type between these two conditions (*e.g*. 313 v. 135 upregulated, **Fig. S3** and **Table S4**). Moreover, 92 (68%) of the 135 infection-upregulated genes in wild type were also upregulated in *nhr-49/PPARA* mutants, and categorized as NHR-49/PPAR-α-independent (**Fig. S3C** and **Table S4**). Examples of these genes included lysozymes *ilys-2, ilys-3*, and *lys-3*, and infection response gene *irg-6* (Troemel et al., 2006).

Additionally, 43 genes were induced by infection in wild type but not in *nhr-49/PPARA* mutants, and thus categorized as NHR-49-dependent (**Fig. S3C** and **Table S4**). Examples included C-type lectin *clec-60*, lysozyme *lys-5*, and, importantly, *fmo-2/FMO5* and *K08C7.4*, two of the six HLH-30/TFEB-independent (**Fig. S3C, Table S3 Fig. 2**) and NHR-49/PPAR-α-dependent genes (**Fig. 3 and Fig. S2**). In contrast, 221 genes were induced by infection only in *nhr-49/PPARA* mutants. These included important host defense transcription factors, such as *cebp-1/CEBP* and *pha-4/FOXA1*, infection response genes *irg-2* and *irg-5*, and C-type lectins *clec-70* and *clec-71* (Bolz et al., 2010; Estes et al., 2010; Irazoqui et al., 2008, 2010a; Pukkila-Worley et al., 2012). Thus, it appeared that *nhr-49/PPARA* loss may be compensated by a large infection-specific response that does not normally occur in wild type animals. However, loss of *nhr-49/PPARA* abrogated the induction of less than one-third of the wild type infection-specific signature, suggesting that *nhr-49/PPARA* makes its important contribution to host defense through the induction of relatively fewer genes than *hlh-30/TFEB*.

### HLH-30/TFEB genetically functions downstream of NHR-49/PPAR-α for host defense

During infection, *nhr-49/PPARA* expression did not change in wild type animals compared to uninfected controls (**Fig. 6A**). Moreover, *nhr-49/PPARA* expression was similar in noninfected wild type and *hlh-30/TFEB* mutants. In contrast, in infected *hlh-30/TFEB* mutants compared with wild type, *nhr-49/PPARA* expression was lower (**Fig. 6A**), indicating that HLH-30/TFEB contributes to *nhr-49/PPARA* expression during infection. Conversely, in *nhr-49/PPARA* null mutants *hlh-30/TFEB* baseline expression was higher than in wild type, yet its induction by infection was abrogated (**Fig. 6B**). This indicated that *nhr-49/PPARA* is required for increased expression of *hlh-30/TFEB* during infection. Moreover, in *nhr-49/PPARA* gain-of-function mutants compared to wild type, both *hlh-30/TFEB* baseline expression and induction were higher (**Fig. 6B**). Considered together, these data suggested that HLH-30/TFEB and NHR-49/PPAR-α contribute to each other’s expression in noninfected and infected animals in different ways.

**Figure 6.**
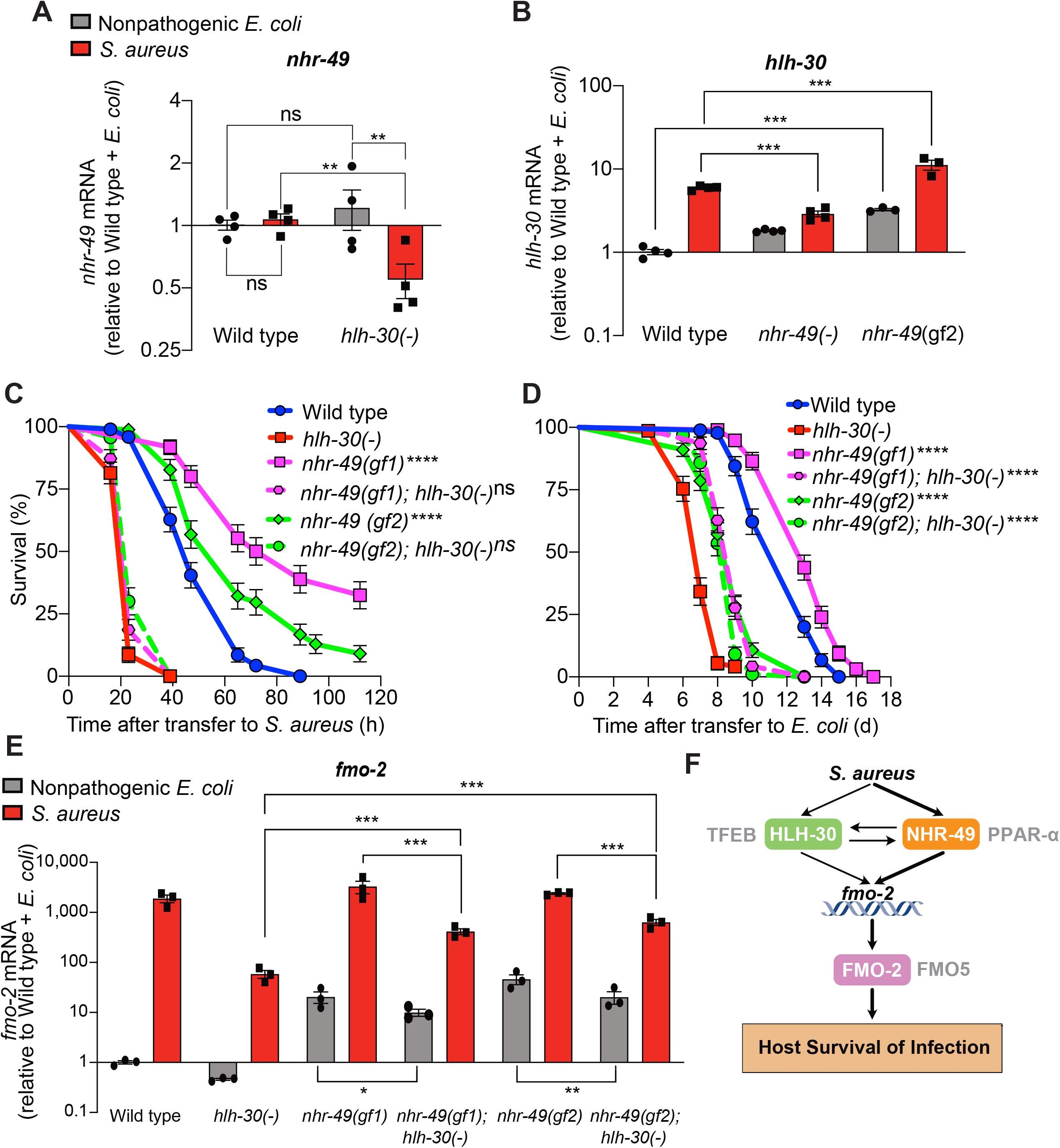
HLH-30/TFEB genetically functions downstream of NHR-49/PPAR-α for host defense. **(A)** Relative expression of *nhr-49/PPARA* transcript (RT-qPCR -ΔCt) in wild type and *hlh-30/TFEB* loss of function mutants fed nonpathogenic *E. coli* OP50 or infected with *S. aureus* SH1000 (8 h). Data are normalized to wild type on *E. coli*, means ± SEM (4 independent biological replicates). ** P < 0.01, ns = not significant, one-way ANOVA followed by Sidak’s test for multiple comparisons. **(B)** Relative expression of *hlh-30/TFEB* transcript (RT-qPCR -ΔCt) in wild type, *nhr-49/PPARA* loss of function, and *nhr-49/PPARA* gain of function (gf2) mutants fed nonpathogenic *E. coli* OP50 or infected with *S. aureus* SH1000 (8 h). Data are normalized to wild type on *E. coli*, means ± SEM (3 - 4 independent biological replicates, indicated). *** P < 0.001, one-way ANOVA followed by Sidak’s test for multiple comparisons. **(C)** Survival of wild type, *hlh-30/TFEB* loss of function, *nhr-49*(*gf1*), *nhr-49*(*gf2*), *nhr-49(gf1); hlh-30(-)*, and *nhr-49(gf2); hlh-30(-)* animals infected with *S. aureus* SH1000. Data are representative of 2 independent replicates. **** P < 0.0001 (Log-Rank test, compared to *hlh-30*(-) mutants). **(D)** Lifespan of wild type, *hlh-30(-), nhr-49(gf1), nhr-49(gf2), nhr-49(gf1); hlh-30(-)*, and *nhr-49(gf2); hlh-30(-)* animals on nonpathogenic *E. coli* OP50. Data are representative of 2 independent replicates. ““ P < 0.0001 (Log-Rank test, compared to *hlh-30*(-) mutants). **(E)** Relative expression of *fmo-2/FMO5* transcript (RT-qPCR -ΔCt) in wild type, *hlh-30(-), nhr-49(gf1), nhr-49(gf1);hlh-30(-), nhr-49(gf2)*, and *nhr-49(gf2);hlh-30*(-) animals fed nonpathogenic *E. coli* OP50 or infected with *S. aureus* SH1000 (4 h). Data are normalized to wild type on *E. coli*, means ± SEM (3 - 4 independent biological replicates). * P ≤ 0.05, ** P < 0.01, *** P < 0.001, one-way ANOVA followed by Sidak’s test for multiple comparisons. **(F)** Schematic representation of *fmo-2/FMO5* regulation during infection with *S. aureus*. Human homologs of the *C. elegans* proteins are indicated in grey lettering.

Because *hlh-30/TFEB* and *nhr-49/PPARA* contributed to both infection-specific *fmo-2/FMO5* induction (**Fig. 2, Fig. 3**) and each other’s expression (**Fig. 6A, B**), we examined their genetic interactions. To determine whether *hlh-30-30/TFEB* and *nhr-49/PPARA* genetically function in the same pathway, we attempted to generate *hlh-30*(-); *nhr-49*(-) double mutants, but were unable to obtain them from genetic crosses suggesting synthetic lethality, consistent with a previous report (Goh et al., 2018). However, it was possible to construct *nhr-49/PPARA* (gain-of-function); *hlh-30/TFEB* (loss-of-function) double mutants. Remarkably, neither *nhr-49*(*gf1*) nor *nhr-49*(gf2), the two mutations that caused enhanced infection survival (**Fig. 4C**), rescued the diminished infection survival of *hlh-30/TFEB* mutants (**Fig. 6C**). In contrast, lifespan on nonpathogenic *E. coli* was increased in both cases, as compared to *hlh-30/TFEB* single mutants (**Fig. 6D**). These data showed that during infection *hlh-30/TFEB* is epistatic to *nhr-49/PPARA*, suggesting that *hlh-30/TFEB* genetically functions downstream of *nhr-49/PPARA* for infection survival, as suggested by *hlh-30/TFEB* expression in *nhr-49/PPARA* mutants (**Fig. 6B**). For longevity, the effect of *nhr-49/PPARA* gain of function and *hlh-30/TFEB* loss of function was additive, suggesting that they regulate aging in parallel genetic pathways.

Expression of *fmo-2/FMO5* mirrored these genetic interactions (**Fig. 6E**). In noninfected animals, incorporation of *hlh-30* (loss of function) mildly affected the high constitutive expression of *fmo-2/FMO5* in *nhr-49/PPARA* (gain of function) mutants (**Fig. 6E**). In contrast, infected *nhr-49/PPARA* (gain of function); *hlh-30/TFEB* (loss of function) double mutants exhibited the *hlh-30/TFEB* (loss of function) phenotype, *i.e*. decreased *fmo-2/FMO5* expression compared to *nhr-49/PPARA* (gain of function, gf1 and gf2) (**Fig. 6E**). Thus, *hlh-30/TFEB* was epistatic to *nhr-49/PPARA* for *fmo-2/FMO5* expression during infection, consistent with it acting downstream or parallel to *nhr-49/PPARA* for infection survival.

### FMO-2/FMO5 is required for host survival of infection

So far, we focused on *fmo-2/FMO5* as a useful reporter of the host response, but its biological relevance to infection survival was unclear. FMO-2 and FMO5 belong to the evolutionarily-conserved flavin-containing monooxygenase (FMO) protein family (Huijbers et al., 2014). In mammals, FMO proteins are primarily known to function in the detoxification of foreign substances (xenobiotics) with prominent roles in drug metabolism (Krueger and Williams, 2005). *C. elegans* FMO-2 exhibits homology to human proteins FMO1-5, with closest similarity to FMO5 (42% identity). Previously, FMO-2/FMO5 had been implicated in dietary-restriction-mediated lifespan extension, and its forced expression resulted in stress resistance (Leiser et al., 2015). In plants, FMOs participate in host defense against bacterial and fungal infections (Bartsch et al., 2006; Koch et al., 2006). Whether animal FMOs also function in innate host defense was not known.

To determine the physiological relevance of *fmo-2/FMO5* during infection, we examined mutants homozygous for a deletion in *fmo-2/FMO5* predicted to result in a null allele (C. elegans Deletion Mutant Consortium, 2012). Compared with wild type, *fmo-2/FMO5* mutants exhibited greatly compromised survival of *S. aureus* infection (**Fig. 7A**) but did not exhibit differences in survival of *P. aeruginosa* infection or in aging when fed nonpathogenic *E. coli* (**Fig. 7B-C**). Deletion of *fmo-2/FMO5* did not affect the induction of the 9 most highly induced infection-specific signature genes (**Fig. S4**). These data suggested that FMO-2/FMO5 may play an important role for host defense specifically during *S. aureus* infection, which is independent of the induction of many other host defense genes.

**Figure 7.**
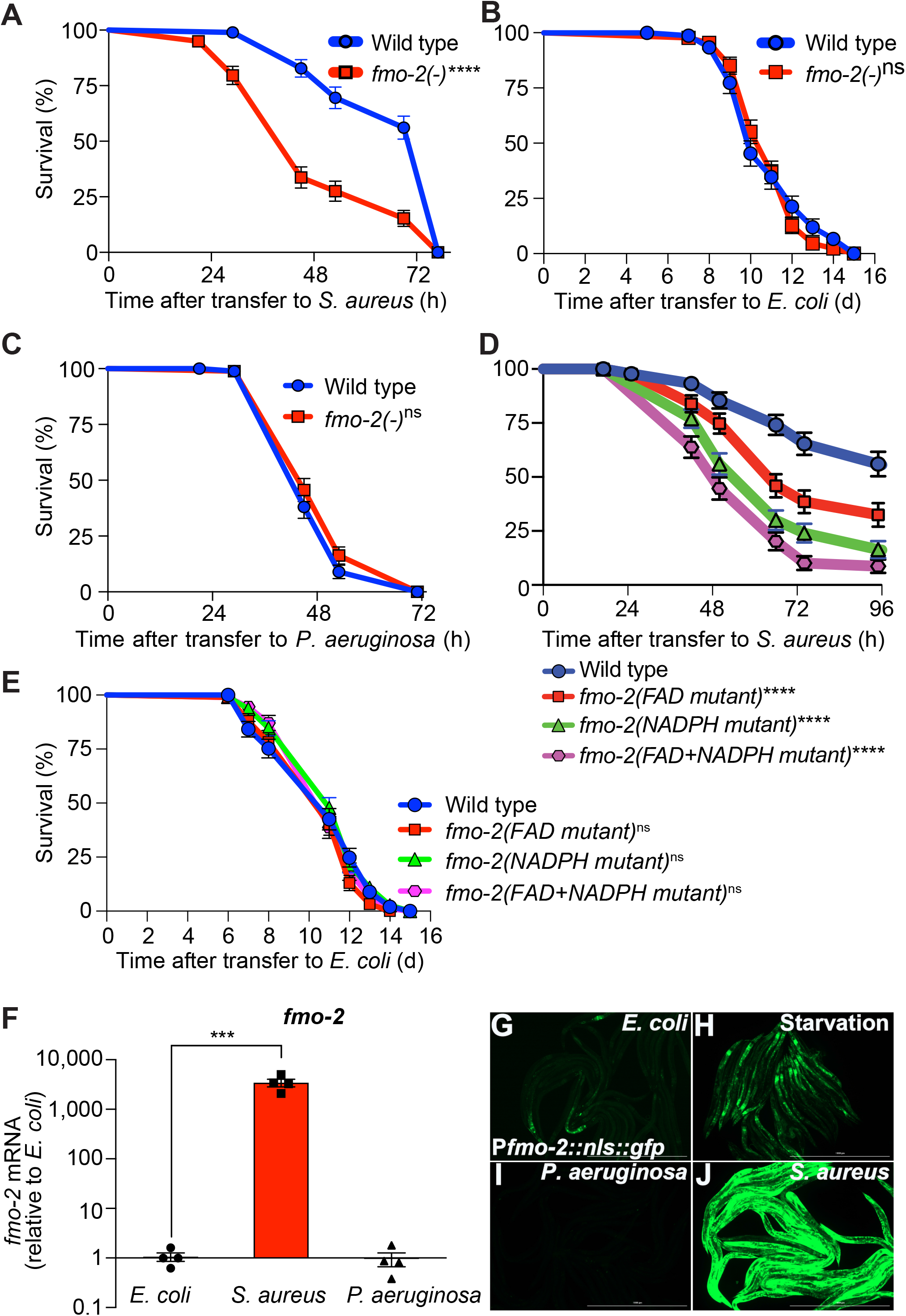
FMO-2/FMO5 is required for host survival of infection. **(A)** Survival of wild type and *fmo-2/FMO5* loss of function mutants infected with *S. aureus* SH1000. Data are representative of 5 independent replicates. **** P < 0.0001 (Log-Rank test). **(B)** Lifespan of wild type and *fmo-2/FMO5* loss of function mutants fed nonpathogenic *E. coli* OP50. Data are representative of 3 independent replicates. ns = not significant (Log-Rank test). **(C)** Survival of wild type and *fmo-2/FMO5* loss of function mutants infected with *P. aeruginosa* PA14. Data are representative of 2 independent replicates. ns = not significant (Log-Rank test). **(D)** Survival of wild type and *fmo-2(FAD), fmo-2(NADPH)*, and *fmo-2(FAD+NADPH)* mutants infected with *S. aureus* SH1000. Data are representative of 2 independent replicates. ****P < 0.0001 (Log-Rank test). **(E)** Lifespan of wild type and *fmo-2(FAD), fmo-2(NADPH)*, and *fmo-2(FAD+NADPH)* mutants on *E. coli* OP50. Data are representative of 3 independent replicates. ns = not significant (Log-Rank test). **(F)** RT-qPCR of *fmo-2/FMO5* transcript in wild type animals fed nonpathogenic *E. coli*, or infected with *S. aureus* SH1000 or *P. aeruginosa* PA14 for 4 h. Data are normalized to *E. coli*, means ± SEM (4 independent biological replicates). *** P ≤ 0.001, unpaired two-sample two-tailed *t*-test. **(G-J)** Epifluorescence micrographs of animals expressing NLS-GFP driven by the endogenous *fmo-2/FMO5* promoter (P*fmo-2::nls::gfp*) fed on *E. coli*, infected with *S. aureus* SH1000 or *P. aeruginosa* PA14, or starved (4 h). Scale bar = 1,000 μm.

To determine whether such a role of FMO-2/FMO5 requires its catalytic activity, we used CRISPR-mediated genome editing to modify key conserved residues in the FMO-2/FMO5 FAD-binding domain, the NADPH-binding domain, or both (**Fig. S5A-B)**. Due to their conservation in FMOs from yeast, plants, and animals **(Fig. S5B)**, these residues are predicted to be required for electron transfer from organic substrates to cofactors FAD and NADPH (Kubo et al., 1997; Rescigno and Perham, 1994). Remarkably, mutation of the NADPH binding site caused a severe infection survival defect, while mutation of the FAD binding site caused a somewhat milder phenotype (**Fig. 7D**). Mutation of both binding sites produced an additive defect (**Fig. 7D**). These results suggested that both cofactor binding sites were required for FMO-2/FMO5 function in host defense. In contrast, none of these mutations, alone or in combination, altered total lifespan on nonpathogenic *E. coli* (**Fig. 7E**). Together, these data indicated that FMO-2/FMO5 catalytic activity may be specifically required for host defense against infection.

Simultaneous deletion of *nhr-49/PPARA* and *fmo-2/FMO5* resulted in an infection survival phenotype that was similar to those of the single mutants (**Fig. S6A**), suggesting that *nhr-49/PPARA* and *fmo-2/FMO5* function in the same genetic pathway. However, the *nhr-49/PPARA* mutant lifespan defect was epistatic to the lack of effect of *fmo-2/FMO5* mutation (**Fig. S6B**), suggesting that *nhr-49/PPARA* may regulate aging independently of *fmo-2/FMO5*.

As mentioned, *fmo-2/FMO5* transcript was induced several thousand-fold in *S. aureus-*infected animals relative to nonpathogenic *E. coli* controls (**Fig. 7F**). In contrast, animals infected with Gram-negative pathogenic bacterium *Pseudomonas aeruginosa* exhibited no significant change (**Fig. 7F**), consistent with previous results (Irazoqui et al., 2008, 2010a; Wong et al., 2007). The fluorescent *in vivo fmo-2/FMO5* transcriptional reporter showed faint GFP expression, mostly in the anterior intestine and head of noninfected animals (**Fig. 7G**). Starvation modestly increased GFP expression in the intestine and nervous system (**Fig. 7H**), while *P. aeruginosa* repressed it below the levels observed in noninfected animals (**Fig. 7I**). In stark contrast, *S. aureus* caused high GFP induction in all tissues, except in gonads and eggs (**Fig. 7J**). These observations confirmed that *fmo-2/FMO5* is strongly induced in a pathogen-specific manner.

Moreover, we found that intestinal-restricted *fmo-2/FMO5* overexpression was sufficient to boost infection survival (**Fig. S7A**). Interestingly, the lifespan of these animals was also extended on nonpathogenic *E. coli* (**Fig. S7B**) in accordance with previous reports (Leiser et al., 2015). These results suggested that elevating FMO-2/FMO5 levels in the intestine alone confers benefits not only in host defense but also against aging, possibly by increasing host resistance to food *E. coli* pathogenesis late in life (McGee et al., 2011; Zhao et al., 2017). Altogether, these observations suggested that *fmo-2/FMO5* is necessary and sufficient for host defense against *S. aureus*.

## DISCUSSION

Because bacteria serve as nutritional source for *C. elegans*, and because intestinal infections cause destruction of the epithelium resulting in loss of nutrient absorption, transcriptional responses to nutritional challenges are likely intertwined with the transcriptional host defense response to the pathogen itself. This raises the question of whether *C. elegans* senses infection as a stress *per se*, through its physiological consequences in the organism, or a combination of both. Here, by directly comparing transcriptomes of animals that were infected with *S. aureus* or were starved, we discovered that starvation and infection elicit large and distinct transcriptional signatures. This indicates that the *C. elegans* host response to *S. aureus* infection is not entirely the result of starvation, and enables the dissection of infection-specific and starvation-specific host response regulatory modules as shown here.

In the present study, we found that loss of HLH-30/TFEB almost completely abrogated differential gene expression between starvation and infection – implicating HLH-30/TFEB not just in a hypothetical overlapping response but in each of these two distinct signatures. This strongly suggests that HLH-30/TFEB integrates metabolic and other stresses to contribute to stress-specific transcriptional responses. The molecular mechanisms that enable a single transcription factor to mediate specific transcriptional responses to distinct stresses may involve stress-specific signals or transcriptional co-factors.

By focusing on *fmo-2/FMO5*, which is highly and specifically induced by infection and is only partially dependent on HLH-30/TFEB, we discovered a novel role for the nuclear receptor NHR-49/PPAR-α in host defense against infection. NHR-49/PPAR-α is better known in *C. elegans* as a transcription factor that is important for the response to starvation (Van Gilst et al., 2005). However, recently NHR-49/PPAR-α was shown to mediate the defense response to exogenous oxidative stress (Goh et al., 2018; Hu et al., 2018). Thus, NHR-49/PPAR-α participates in host defense against biotic and abiotic stressors, and should be considered a key player in the organismal stress response alongside SKN-1/NRF, DAF-16/FOXO3, and HLH-30/TFEB (Blackwell et al., 2015; Lin et al., 2018; Tissenbaum, 2018). Our analysis showed that NHR-49/PPAR-α is not required for as large a portion of the host response to infection as HLH-30/TFEB, even though NHR-49/PPAR-α is partially required for HLH-30/TFEB induction. The larger HLH-30/TFEB regulon implies that during infection signals in addition to NHR-49/PPAR-α activation contribute to HLH-30/TFEB regulation. Similar to HLH-30/TFEB, how NHR-49/PPAR-α induces specific responses to distinct stresses is also unknown. These findings are relevant beyond nematodes, as PPAR-α regulates TFEB in mammalian cells (Kim et al., 2017). Moreover, the microbiota represses HNF4-α, a second NHR-49 homolog, in zebrafish and mice, to maintain intestinal homeostasis (Davison et al., 2017). Therefore, unraveling the control of NHR-49/PPAR-α in relation to intestinal microbiota and infection may provide useful information to understand vertebrate intestinal homeostasis and host defense.

In addition to leading us to discover NHR-49/PPAR-α, FMO-2/FMO5 is interesting in its own right. Regulation of *fmo-2/FMO5* is complex. During infection, NHR-49/PPAR-α appeared to drive *fmo-2/FMO5* expression in the pharynx, nervous system, and anterior intestine, while *hlh-30/TFEB* induced it in the intestine, muscle, and epidermis in complementary spatial patterns. Hypoxia and dietary restriction induce *fmo-2/FMO5* (Leiser et al., 2015; Shen et al., 2005), and so do gain of function mutations in *hif-1/HIF1* or *skn-1/NRF2* (Leiser et al., 2015; Nhan et al., 2019). Lifespan extension by dietary restriction or *hif-1/HIF1* gain of function requires *fmo-2/FMO5;* moreover, *hlh-30/TFEB* loss of function quenches lifespan extension by *hif-1/HIF1* and *fmo-2/FMO5* induction by hypoxia and fasting (Leiser et al., 2015). These observations lend further support to our findings that HLH-30/TFEB partially induces *fmo-2/FMO5* during infection.

In addition, we found that loss of FMO-2/FMO5 causes a severe defect in infection survival without affecting longevity. Thus, FMO-2/FMO5 represents a novel host defense effector. We previously examined the requirement for *fmo-2/FMO5* using RNAi-mediated silencing, but such manipulation failed to produce a phenotype for reasons unknown (Irazoqui et al., 2010a). Moreover, the failure of tissue-specific RNAi to elicit a phenotype and the toxicity of *fmo-2/FMO5* extrachromosomal transgenic constructs precluded our investigation of the tissues of *fmo-2/FMO5* action for host defense. However, single copy intestinal expression of FMO-2/FMO5 boosted host defense, suggesting that FMO-2/FMO5 could play a major role in the intestine, a hub for host defense in *C. elegans* (McGhee, 2007). Nonetheless, FMO-2/FMO5 induction appears to be a major mechanism of host defense in *C. elegans*. Exactly how FMO-2/FMO5 promotes host infection survival is poorly understood, but site-directed mutagenesis of the NADPH and FAD binding sites revealed that the mechanism of action requires its catalytic activity. In addition, human FMO5 can generate large amounts of H2O2 from O2 (Fiorentini et al., 2016). Thus, it is possible that FMO-2/FMO5 is an infection-specific NADPH oxidase that generates H_2_O_2_ with antimicrobial and signaling functions (McCallum and Garsin, 2016; Sies and Jones, 2020). The observed roles of *fmo-2/FMO5* in survival of heat, di-thiothreitol, and tunicamycin stress are consistent with a H2O2-mediated signaling role (Leiser et al., 2015).

FMOs are emerging as important host defense factors across phylogeny. In plants, FMO1 is required for the catalysis of pipecolic acid to N-hydroxypipecolic acid, which provides systemic acquired resistance to bacterial and oomycete infections (Hartmann et al., 2018). In mammals, FMO3 is an evolutionarily ancient FMO that exhibits unique substrate specificity and catalyzes multiple drugs that is important for their detoxification (Krueger and Williams, 2005). However, to date no reports have indicated an important role for FMO5, or other FMOs, in mammalian (or any animal) innate immunity. In mice, FMO5 is expressed in many tissues and organs, including the liver and the epithelium of the gastrointestinal tract (Scott et al., 2017). Mouse FMO5 is required for sensing the microbiota, and *Fmo5^-/-^* mutants exhibit altered metabolic profiles and microbiomes compared with wild type mice (Scott et al., 2017). Furthermore, *Fmo5*^-/-^ mutants exhibit a 70% reduction in plasma TNF-α compared with wild type (Scott et al., 2017). Together, these observations suggest that FMO5 is an important microbiota sensor and effector that modulates the intestinal microbiota, but the mechanism of action is unknown. Therefore, elucidation of mechanisms of host defense mediated by *fmo-2* in nematodes and *FMO5* in mammals will provide fundamental insight into evolutionarily conserved mechanisms of host defense against infection and identify therapeutic opportunities for infections and inflammatory diseases.

## ACKNOWLEDGEMENTS

The authors are grateful to members of the Irazoqui laboratory, the Program in Innate Immunity, and the Department of Microbiology and Physiological Systems for helpful insights and discussions. Joyce Barrett, Linda Benson, Richard Fish, Amy Parker, Cheryl Barry, and Tammy Bailey provided expert facilities and administrative assistance. Scott F. Leiser (University of Michigan) provided KAE11 strain. Some strains used in this study were provided by the *Caenorhabditis* Genetics Center, which is funded by the NIH Office of Research Infrastructure Programs (P40-OD010440). Research reported in this publication was supported by the National Institute of General Medical Sciences of the National Institutes of Health under award number GM101056, and by the National Science Foundation under award number NSF1457055 (to J.E.I.), and by a grant from the National Institutes of Aging (R01AG051659) to A.G. The content is solely the responsibility of the authors and does not necessarily represent the official views of the National Institutes of Health.

## AUTHOR CONTRIBUTIONS

K.A.W. and J.E.I. conceived and designed the experiments. K.A.W. and J.E.I. analyzed the data. K.A.W. and D.G. performed the experiments. S.T., R.R., and A.G. provided reagents and essential intellectual input. All authors participated in manuscript writing and editing.

## MATERIALS AND METHODS

### KEY RESORCES TABLE

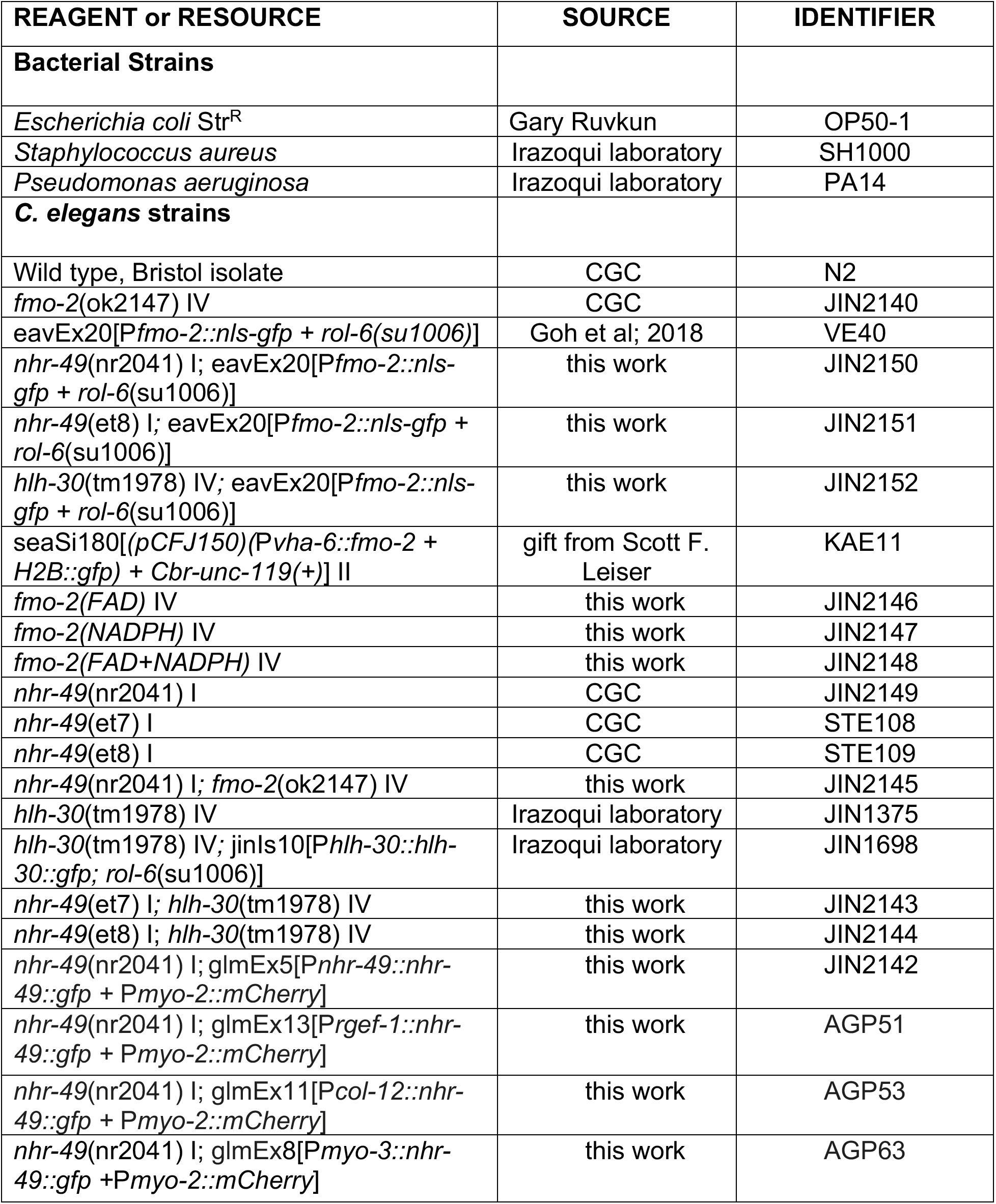
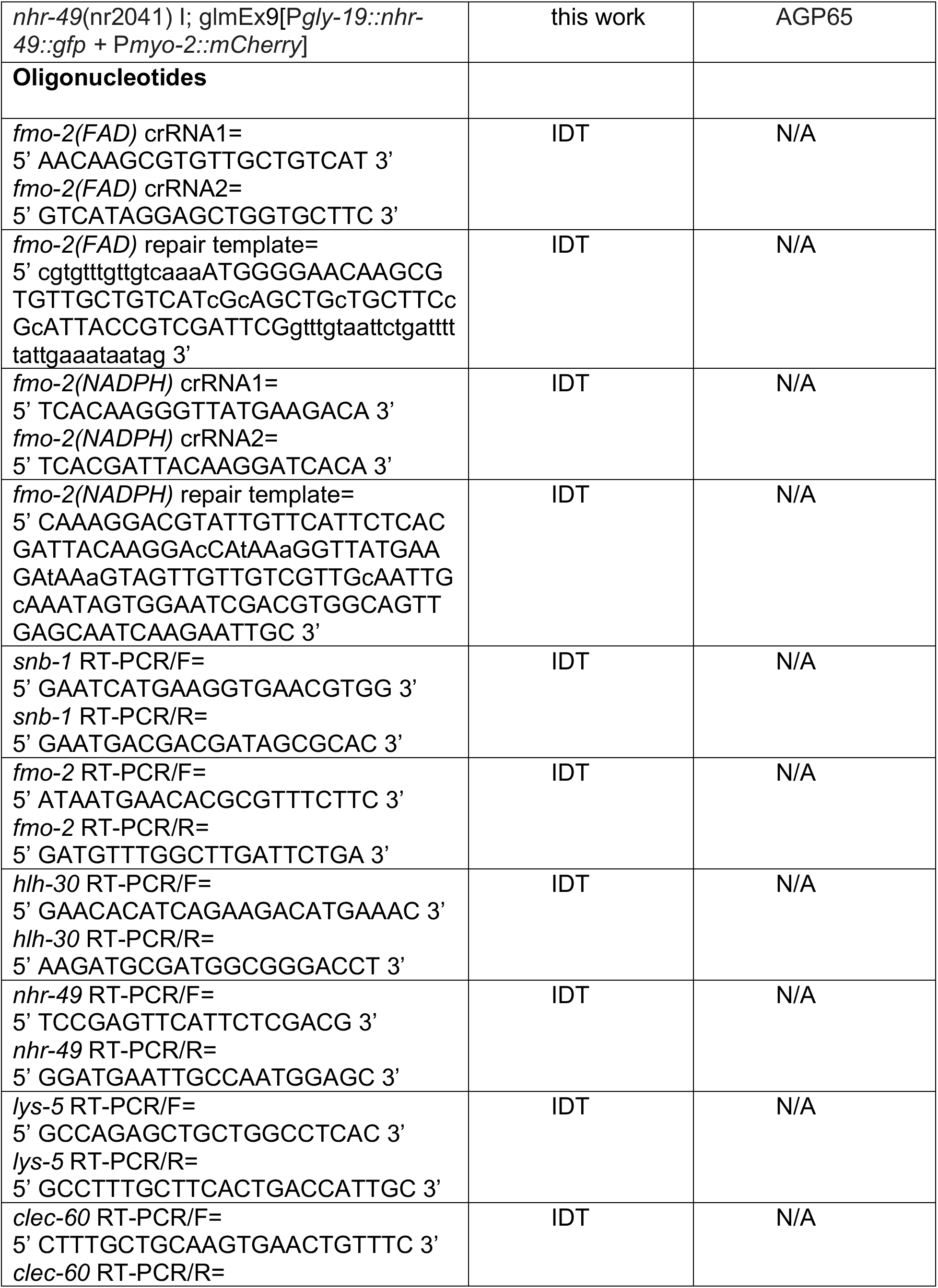
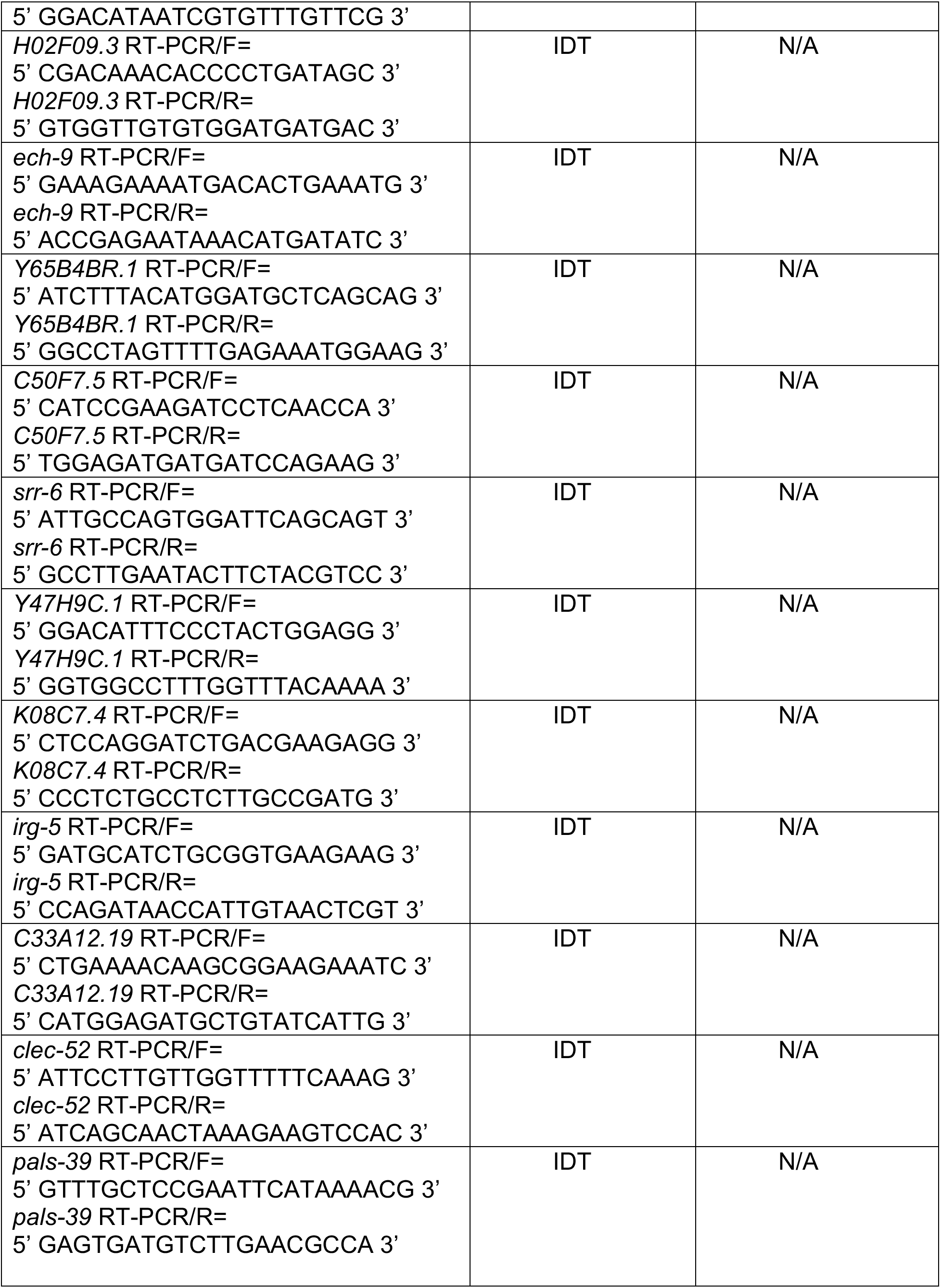
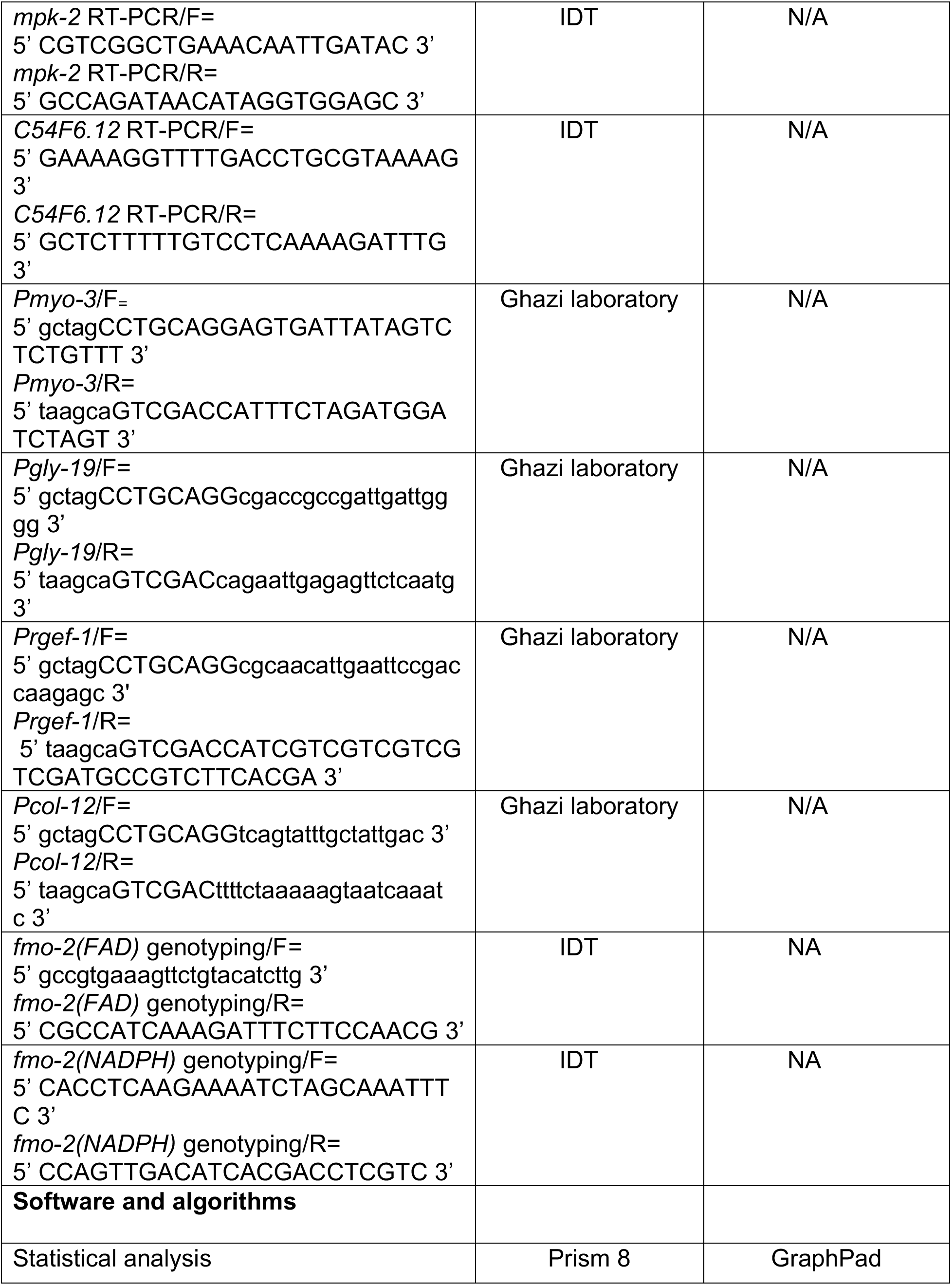
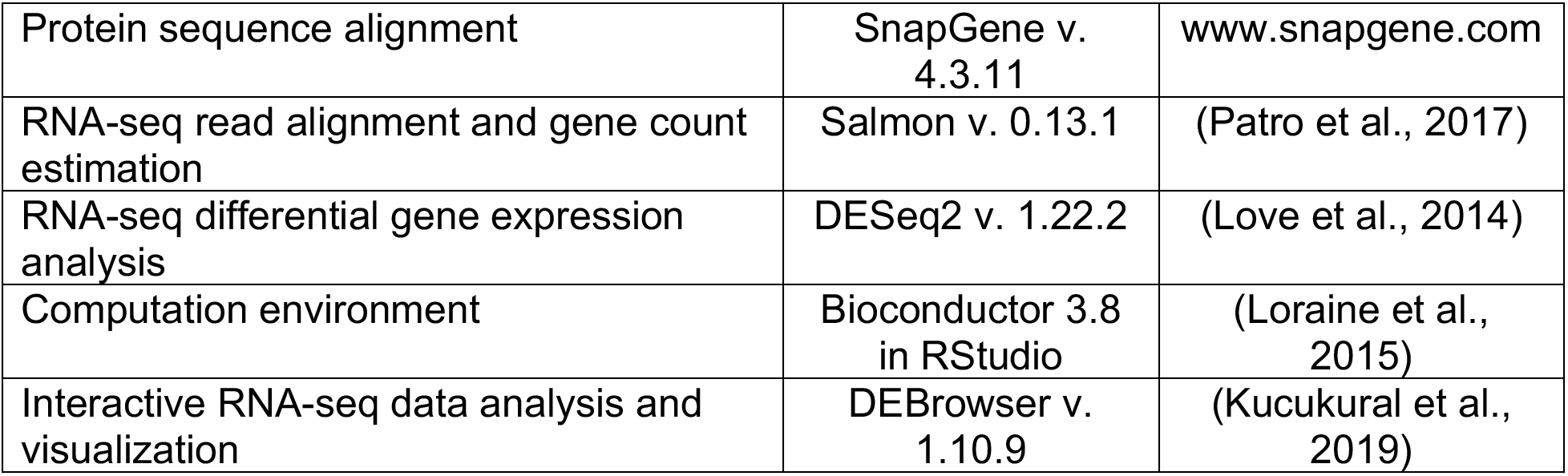

#### Experimental model

The nematode *C. elegans* was used as the experimental model for this study. Strains were maintained at 15 – 20 °C on Nematode Growth Media (NGM) plates seeded with Str^R^ *E. coli* OP50-1 strain using standard methods (Stiernagle, 2006).

#### Method Details

##### Infection assays

*S. aureus* SH1000 strain was grown overnight in tryptic soy broth (TSB) containing 50 μg/ml kanamycin (KAN). Overnight cultures were diluted 1:1 with TSB and 10 μl of the diluted culture was uniformly spread on the entire surface of 35 mm tryptic soy agar (TSA) plates containing 10 μg/ml KAN. Plates were incubated for 5 - 6 h at 37 °C, then stored overnight at 4 °C. *P. aeruginosa* isolate PA14 was grown overnight in Luria broth. 10 μl of the overnight culture was uniformly spread on the entire surface of 35 mm NGM plates. Plates were incubated at 37 °C for 24 h followed by 25 °C for 48 h (Powell and Ausubel, 2008). Animals were treated with 100 μg/ml 5-fluoro-2’-deoxyuridine (FUDR) at L4 larval stage for ~24 h at 15 °C - 20 °C before transfer to *S. aureus* or *P. aeruginosa* plates. Three plates were assayed for each strain in each replicate, with 20 - 40 animals per plate. Survival was quantified using standard methods (Powell and Ausubel, 2008). Animals that crawled off the plate or died of bursting vulva were censored. Infection assays were carried out at least twice.

##### *S. aureus* infection for RNA analysis

To prepare infection plates, *S. aureus* SH1000 was grown overnight in TSB containing 50 μg/ml KAN. 500 – 1,000 μl of overnight culture was uniformly spread on the entire surface of freshly prepared 100 mm TSA plates supplemented with 10 μg/ml KAN. The plates were incubated for 6 h at 37 °C, then stored overnight at 4 °C. To prepare *P. aeruginosa* plates, *P. aeruginosa* isolate PA14 was grown overnight in Luria broth. 1 ml of overnight culture was uniformly spread on the entire surface of freshly prepared 100 mm NGM plates. The plates were first incubated at 37 °C for 24 h and then at 25 °C for 48 h. To prepare control plates with nonpathogenic *E. coli*, 1 ml of 10 - 20X concentrated overnight culture of OP50-1 bacteria was spread on 100 mm NGM plates, incubated for 5 – 6 h at 37 °C, and then stored at 4 °C, similar to *S. aureus* plates. To prepare plates for starvation, TSA plates were treated similarly to infection plates, except that nothing was added to them. Synchronized young adults of wild type and mutants were seeded the next day on *S. aureus, P. aeruginosa*, OP50-1, and starvation plates that were previously warmed to room temperature. After 4 h incubation at 25 °C, animals for all conditions were washed 3 - 4 times in water, and then lysed in 1 ml of TRIzol reagent (Invitrogen). The samples were snap frozen in liquid nitrogen, then stored at −80 °C. RNA was extracted using 1-bromo-3-chloropropane (MRC) followed by purification with isopropanol-ethanol precipitation. RNA was analyzed by qPCR or sequencing. For sequencing, RNA was additionally purified using PureLink^™^ RNA Mini Kit (Invitrogen). Four independent biological replicates were submitted to BGI for library preparation and sequencing using BGI-seq 500.

##### Longevity (aging) assays

Animals were transferred to 60 mm NGM plates seeded with 10 - 20X concentrated *E. coli* OP50-1 bacteria supplemented with 100 μg/ml FUDR. For consistency with infection assays, longevity assays were also performed at 25 °C. Three plates were assayed for each strain in each replicate, with 20 - 40 animals per plate. Experiments were repeated at least twice. Animals that did not respond to prodding were scored as dead, and the animals that died from bursting vulva or crawled off the plate were censored.

##### Quantitative RT-PCR

After each treatment, *C. elegans* were collected in sterile water and lysed using TRIzol Reagent (Invitrogen). Total RNA was extracted and purified as described before and then digested with DNAse (Bio-Rad). 100 - 1,000 ng of total RNA was used for cDNA synthesis using iScript cDNA synthesis kit (Bio-Rad). RT-qPCR was performed using SYBR Green Supermix (Bio-Rad) using a ViiA7 Real-Time qPCR system (Applied Biosystems). Primer sequences are provided in **Key Resources Table**. At least two independent biological replicates were used for each treatment and *C. elegans* strain. qPCR Ct values were normalized to the *snb-1* control gene, which did not change with the conditions tested, to calculate RT-qPCR ΔCt values. Data analysis was carried out using the Pfaffl method (Pfaffl, 2001). Heat maps were generated using open access online tool Morpheus (https://software.broadinstitute.org/morpheus).

##### Generation of transgenic strains

To construct *Pnhr-49::nhr-49::gfp* containing plasmid, a 6.6 kb genomic fragment of *nhr-49* gene (comprising of 4.4 kb coding region covering all *nhr-49* transcripts plus 2.2 kb sequence upstream of ATG) was cloned into the GFP expression vector pPD95.77 (Addgene #1495), as reported previously (Ratnappan et al., 2014). For generating tissue-specific constructs, the *nhr-49* promoter was replaced with tissue-specific promoters using *Sbf*I and *Sal*I restriction enzymes. The primers that were used to amplify tissue-specific promoters are listed in **Key Resources Table**. For the generation of rescue strains, each rescue plasmid (100 ng/μl) was injected along with pharyngeal muscle-specific *Pmyo-2::mCherry* co-injection marker (25 ng/μl) into *nhr-49*(*nr2041*) mutant strain, using standard methods (Mello and Fire, 1995). Strains were maintained by picking animals that were positive for both GFP and mCherry.

*fmo-2*(*FAD*), *fmo-2*(*NADPH*), and *fmo-2*(*FAD+NADPH*) strains were generated using CRISPR-Cas9 genome editing as described (Dokshin et al., 2018). Residues for mutation were selected based on protein sequence alignment and as previously reported (Bartsch et al., 2006). To isolate worms with mutated residue(s) in FAD or NADPH motifs, silent mutations that resulted in restriction enzyme sites (*PvuII* for FAD, and *Ava*II for NADPH) without any change in the amino acid(s) were created in the repair templates. A PCR fragment spanning the mutated nucleotides was amplified from the progeny of the injected worms, followed by digestion with the above-mentioned restriction enzymes. Mutations in FAD and NADPH motifs were confirmed by sequencing PCR fragments amplified from the corresponding regions in the mutant animals. To generate *fmo-2*(*FAD+NADPH*) double mutant, the *fmo-2*(*NADPH*) mutant strain was used as a background for a second round of CRISPR microinjections. Sequences for crRNAs, repair templates, and the genotyping primers used for the construction of these strains are listed in **Key Resources Table.**

##### Image analysis

Images were captured using a Lionheart FX Automatic Microscope (BioTek Instruments) under a 4X objective. 20 - 30 animals were anesthetized using 100 mM NaN3 on a 2% agarose pad immediately prior to imaging. Fluorescence microscopy analysis was independently replicated at least 3 times.

##### RNA sequencing analysis

BGI provided clean reads in FASTQ format. Clean FASTQ files were verified using FastQC (https://www.bioinformatics.babraham.ac.uk/projects/fastqc) using Bioconductor in RStudio (Loraine et al., 2015) and used as input for read mapping in Salmon v.0.9.1 (Patro et al., 2017) using WBCel.235.cdna from Ensembl (www.ensembl.org) as reference transcriptome. Salmon outputs in quant format were used for input in DESeq2 (Love et al., 2014) in Bioconductor in RStudio for count per gene estimation using batch correction. Total counts per gene tables from DESeq2 were used as input for DEBrowser (Kucukural et al., 2019) for verification of transcriptome replicate similarity, data analysis using the built-in DESeq2 algorithm for differential gene expression analysis (adjusted P value ≤ 0.01 was considered significant), visualization, and interactive data mining. Overlap between gene sets was determined using the Venn tool in BioInfoRx (https://bioinforx.com). GO representation analysis was performed using online tool g:Profiler (Raudvere et al., 2019) (https://biit.cs.ut.ee/gprofiler/gost).

##### Quantification and statistical analysis

Prism 8 (GraphPad) was used for statistical analyses. Survival data were compared using the Log-Rank (Mantel-Cox) test. A P value ≤ 0.05 was considered significantly different from control. For comparisons to a single reference, two-sample, two-tailed *t* tests were performed to evaluate differences between ΔCt values (Schmittgen and Livak, 2008). For multiple comparisons, statistical significance was examined by one-way ANOVA followed by Sidak’s post-hoc test. A P value ≤ 0.05 was considered significant.

---

**Figure S1.**
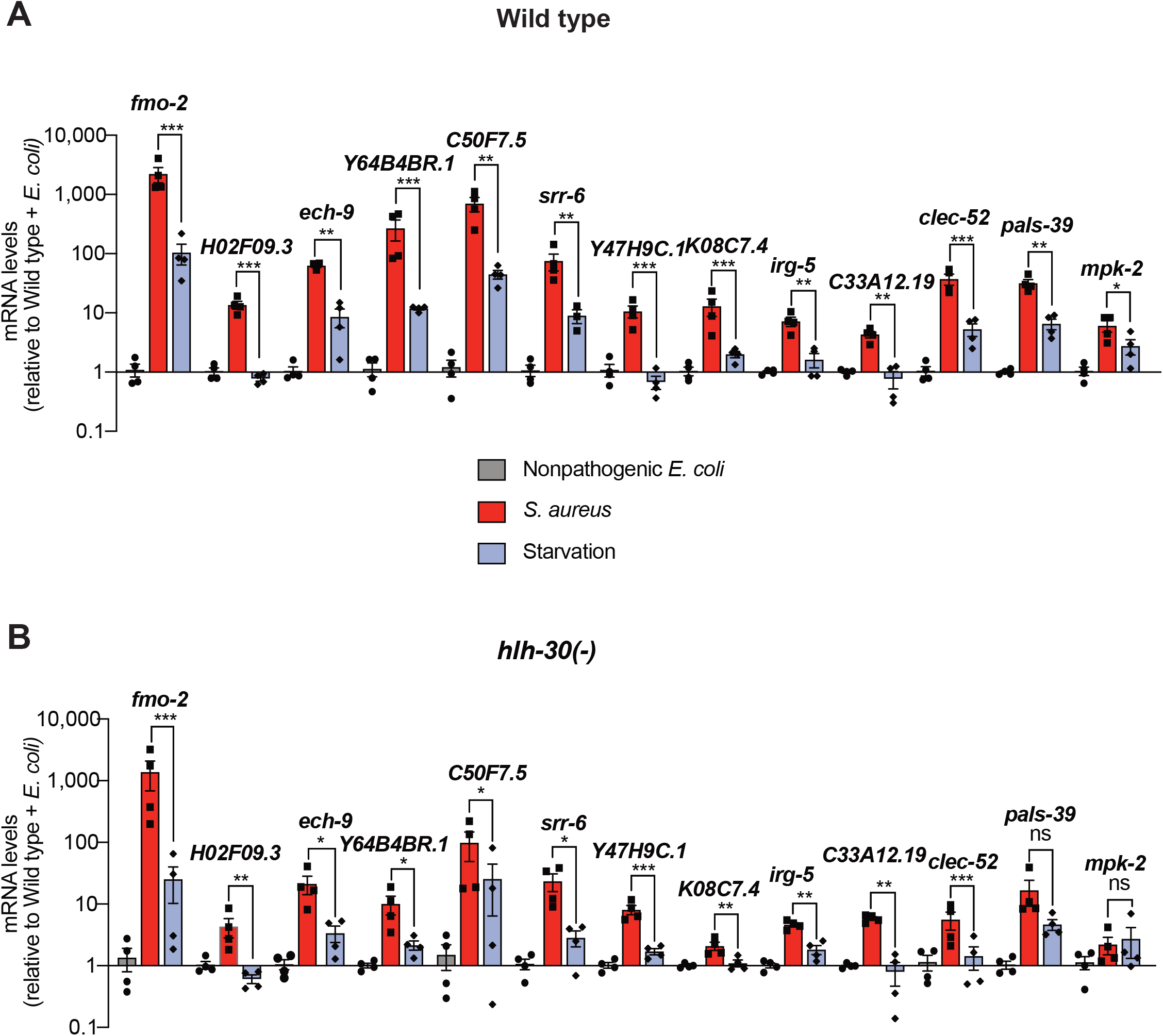
Expression analysis of 13 most highly induced genes (related to Figures 1 and 2). **(A-B)** Relative transcript expression of top most highly induced 13 genes from RNA-seq, measured by RT-qPCR, in wild type and *hlh-30(-)* animals that were fed nonpathogenic *E. coli*, infected with *S. aureus*, or starved for 4 h. Data are normalized to Wild type + *E. coli* and represent mean ± S.E.M. of 3 - 4 biological replicates. ns = not significant, * P < 0.05, ** P < 0.01, *** P < 0.001 (unpaired two-tailed *t* test).

**Figure S2.**
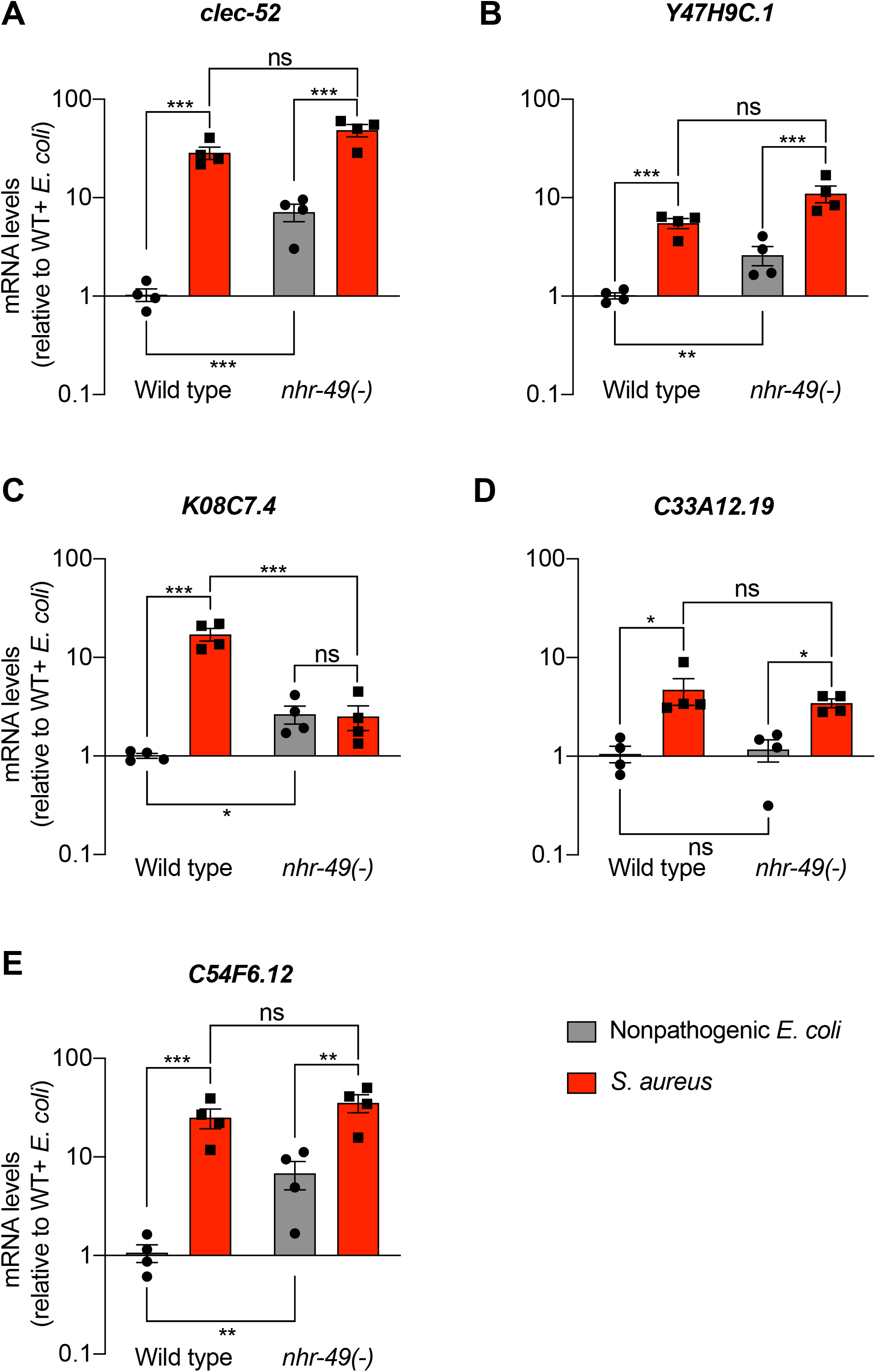
Expression analysis of HLH-30/TFEB-independent genes in *nhr-49/PPARA* mutants (related to Figure 3) Relative transcript expression (RT-qPCR) of 5 genes in wild type and *nhr-49(-)* animals fed nonpathogenic *E. coli* or infected with *S. aureus* (4 h), normalized to wild type + *E. coli*. Data are mean ± SEM (four independent biological replicates). ns = not significant, * P ≤ 0.05; ** P < 0.01; *** P < 0.001, ns = not significant, one-way ANOVA followed by Sidak’s test for multiple comparisons.

**Figure S3.**
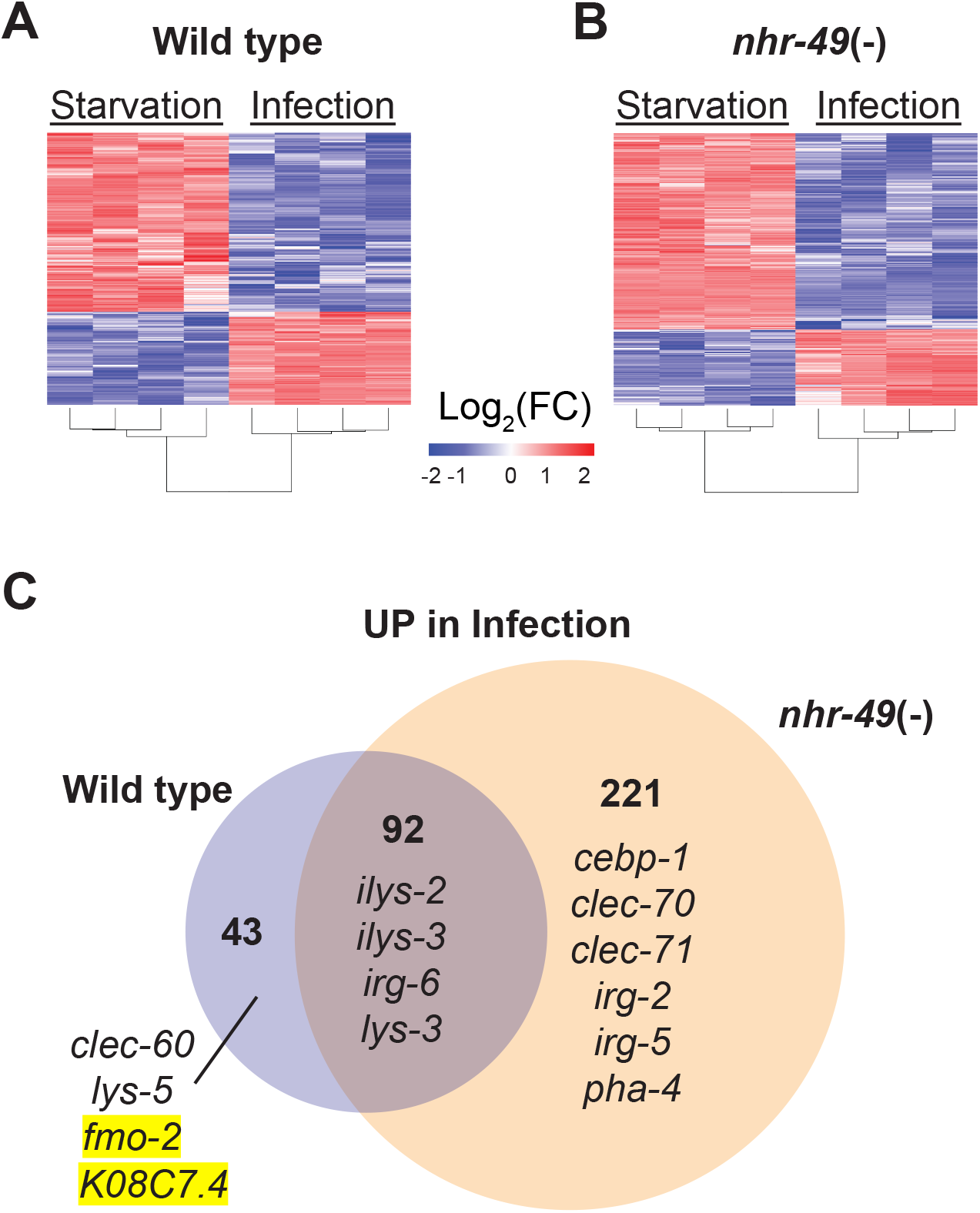
NHR-49/PPAR-α is required for one-third of the infection-specific transcriptional signature. **(A-B)** Heat map of differentially expressed genes during starvation and infection in wild type (A) and *nhr-49(-)* (B) animals (RNA-seq, Log2(FC), PAdj ≤ 0.001). Columns represent a biological replicate each. **(C)** Venn diagram representing genes that are upregulated by 4 h *S. aureus* infection compared with 4 h starvation in wild type and *nhr-49*(-) animals. Shown are a few examples for reference.

**Figure S4.**
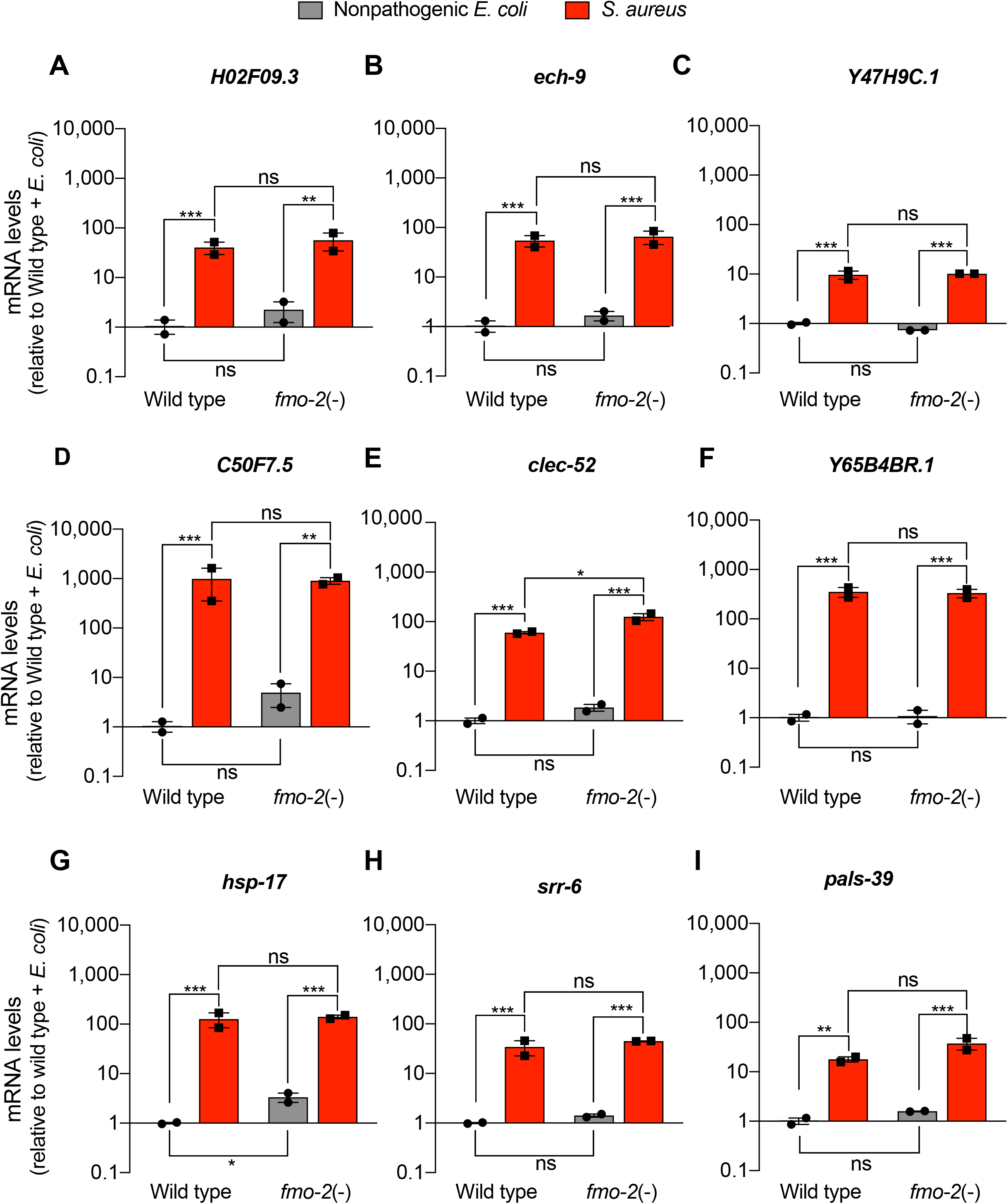
FMO-2/FMO5 is not required for the expression of a set of host defense genes (related to Figure 7). **(A-I)** Relative transcript expression (RT-qPCR) of 9 genes in wild type and *fmo-2(-)* animals fed nonpathogenic *E. coli* or infected with *S. aureus* (4 h), normalized to wild type + *E. coli*. Data are mean ± SEM (two independent biological replicates). ns = not significant, * P ≤ 0.05; ** P < 0.01; *** P < 0.001, ns = not significant, one-way ANOVA followed by Sidak’s test for multiple comparisons.

**Figure S5.**
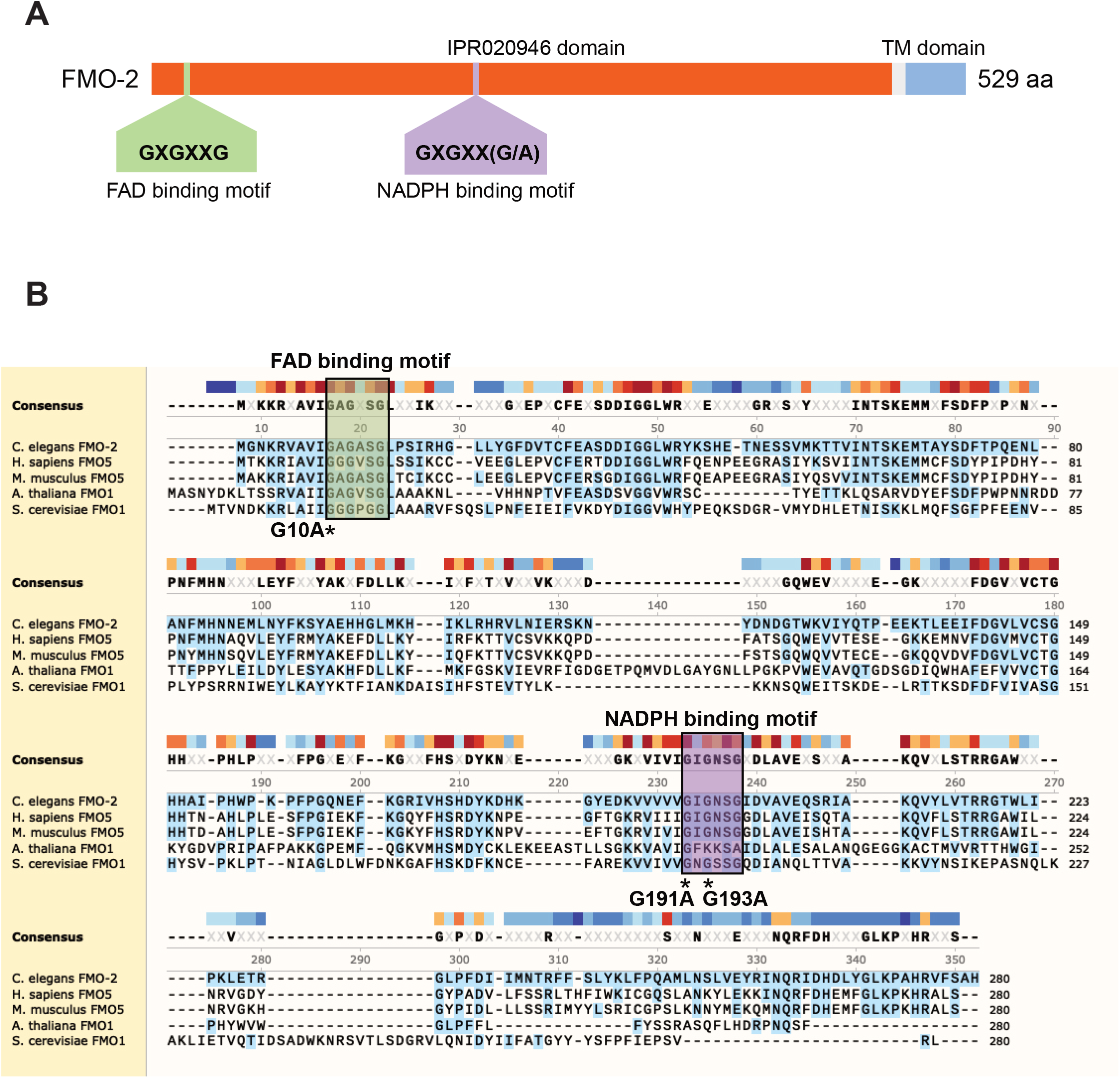
Highly conserved amino acid sequences in FMO-2/FMO5 (related to Figure 7) **(A)** Domain architecture of *C. elegans* FMO-2. Source: InterPro (https://www.ebi.ac.uk/interpro/protein/UniProt/G5EBJ9). TM, transmembrane domain. **(B)** Amino acid sequence alignment of *C. elegans* FMO-2, *Homo sapiens* FMO5, *Mus musculus* FMO5, *Arabidopsis thaliana* FMO1, and *Saccharomyces cerevisiae* Fmo1p. *C. elegans* FMO-2 was used as reference. Protein sequence of up to 280 amino acids was used for alignment in each case. Regions of the proteins that include FAD and NADPH motifs (highlighted in boxes) were chosen to show conservation. Glycine (G) residues modified by CRISPR are indicated with asterisks (*) at the bottom of the amino acids.

**Figure S6.**
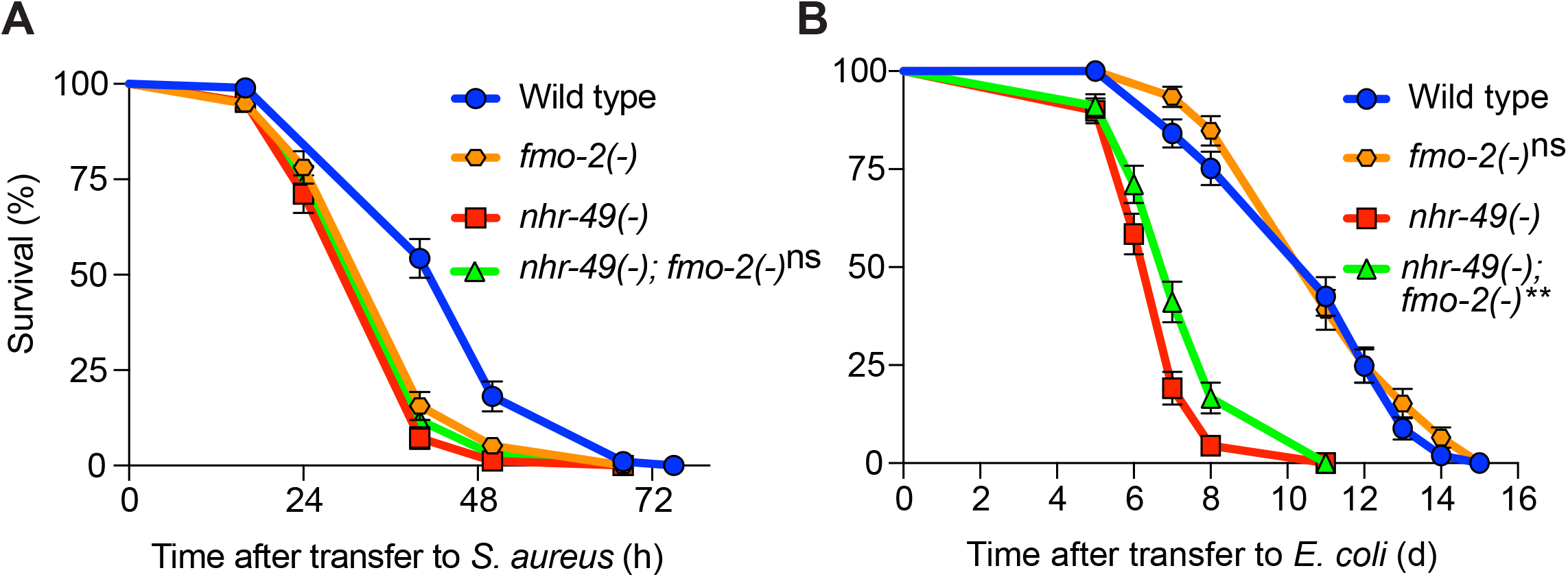
FMO-2/FMO5 and NHR-49/PPAR-α function in the same genetic pathway (related to Figure 7). **(A)** Survival of wild type, *nhr-49(-)*, and *nhr-49(-);fmo-2(-)* animals infected with *S. aureus*. Data are representative of 2 independent replicates. ns = not significant (Log-Rank test). **(B)** Lifespan of wild type, *nhr-49(-)*, and *nhr-49(-)*;*fmo-2(-)* animals on nonpathogenic *E. coli*. Data are representative of 2 independent replicates. ns = not significant, ** P < 0.01 (Log-Rank test).

**Figure S7.**
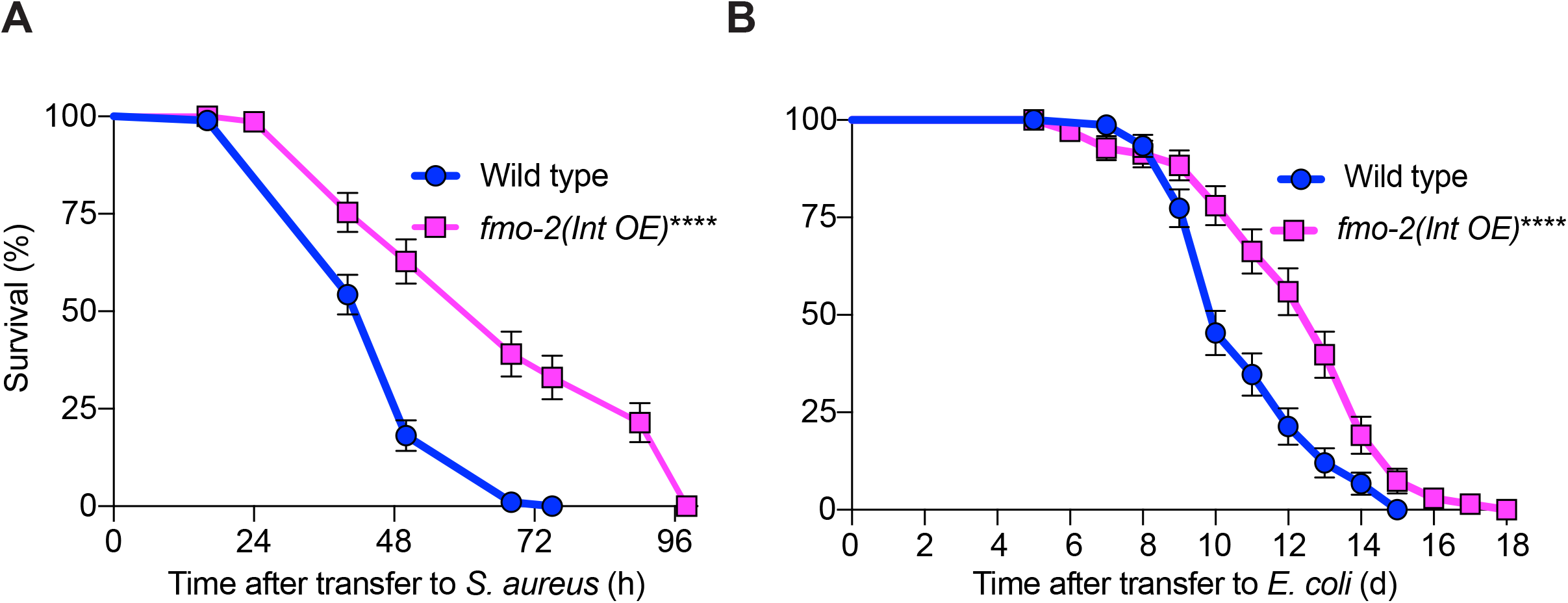
Intestinal overexpression of FMO-2/FMO5 boosts host survival of *S. aureus* infection (related to Figure 7). **(A)** Survival of wild type and intestinal overexpression (OE) line of *fmo-2/FMO5* infected with *S. aureus*. Data are representative of 2 independent replicates. ““ P < 0.0001 (Log-Rank test). Int., intestinal. **(B)** Lifespan of wild type and intestinal overexpression (OE) of *fmo-2/FMO5* on *E. coli* OP50. Data are representative of 2 independent replicates. **** P < 0.0001 (Log-Rank test).

